# Two snakebite antivenoms have potential to reduce Eswatini’s dependency upon a single, increasingly unavailable product: results of preclinical efficacy testing

**DOI:** 10.1101/2022.05.16.492230

**Authors:** Stefanie K. Menzies, Thea Litschka-Koen, Rebecca J. Edge, Jaffer Alsolaiss, Edouard Crittenden, Steven R. Hall, Adam Westhorpe, Brent Thomas, James Murray, Nondusimo Shongwe, Sara Padidar, David G. Lalloo, Nicholas R. Casewell, Jonathan Pons, Robert A. Harrison

## Abstract

**Background:** Snakebite is a major public health concern in Eswatini, where treatment relies upon one antivenom – SAIMR Polyvalent. Although effective in treating snakebite, SAIMR Polyvalent is difficult to source outside its manufacturing country (South Africa) and is dauntingly expensive. We compared the preclinical venom-neutralising efficacy of two alternative antivenoms and SAIMR Polyvalent against the lethal and tissue-destructive effects of venoms from five species of medically important snakes using *in vivo* murine assays. The test antivenoms were ‘Panafrican’ manufactured by Instituto Clodomiro Picado and ‘PANAF’ manufactured by Premium Serums & Vaccines.

**Principal Findings:** *In vivo* murine preclinical studies identified both test antivenoms were equally or more effective than SAIMR Polyvalent at neutralising lethal and tissue-destructive effects of *Naja mossambica* venom. Both test antivenoms were less effective than SAIMR Polyvalent at neutralising the lethal effects of *Bitis arietans, Dendroaspis polylepis, Hemachatus haemachatus* and *Naja annulifera* venoms, but similarly effective at neutralising tissue damage induced by *B. arietans* and *H. haemachatus* venoms.

*In vitro* immunological assays identified that IgG titres and toxin-specificities of the test antivenoms were comparable to SAIMR Polyvalent. Plasma clotting disturbances by *H. haemachatus* and *N. mossambica* were effectively neutralised by the test antivenoms, whereas SAIMR Polyvalent failed to neutralise this bioactivity of *N. mossambica* venom. The SVMP activity of *B. arietans* venom was equally reduced by all three antivenoms. The PLA_2_ activities of *H. haemachatus* and *N. mossambica* were effectively neutralised by all three antivenoms.

**Conclusions:** Panafrican outperformed PANAF, though both were less poly-specifically effective than SAIMR Polyvalent. The efficacy of these antivenoms against the lethal and tissue-destructive effects of *N. mossambica* venom, the most common biting species in Eswatini, identify that Panafrican and PANAF antivenoms offer effective alternatives to SAIMR for the treatment of snakebite in Eswatini, and potentially for neighbouring countries.

**Author Summary:** Treatment of snakebite in Eswatini is reliant upon a single antivenom (SAIMR Polyvalent) manufactured in South Africa. This highly effective product is increasingly difficult to source and is expensive – alternative/additional antivenoms are urgently required to improve patient outcomes following snake envenoming. Using murine preclinical venom toxicity and antivenom efficacy assays, we identified two alternative antivenoms whose venom-neutralising characteristics, while less poly-specifically effective than SAIMR Polyvalent, were as effective against the lethal and tissue-destructive effects of the most common biting snake in Eswatini – *Naja mossambica*. This murine data, already shared with the Eswatini Ministry of Health, supports and justifies human testing of these two antivenoms in Eswatini because increasing the availability of effective and affordable treatments could resolve the current medical dependency of Eswatini snakebite patients upon a single, rarely available and expensive product.

## 1. Introduction

An estimated 138,000 people die annually as a consequence of snakebite, of which 32,000 deaths are estimated to occur in sub-Saharan Africa (sSA) (1). Antivenom is the first choice treatment of snake envenoming and comprises immunoglobulins (Igs) purified from horses or sheep hyperimmunised with snake venoms. The SAIMR Polyvalent antivenom is manufactured in South Africa by South African Vaccine Producers (SAVP) from Igs of horses hyperimmunised with venoms from the ten most medically important snake species found in South Africa and neighbouring countries (see Table 1). The clinical effectiveness of SAIMR Polyvalent in treating snakebite in sSA is widely recognised and reported (2–4). Perhaps because SAIMR Polyvalent is such a highly regarded antivenom (3), South Africa and other southern African countries, including Eswatini, are entirely reliant upon this single product.

**Table 1.**
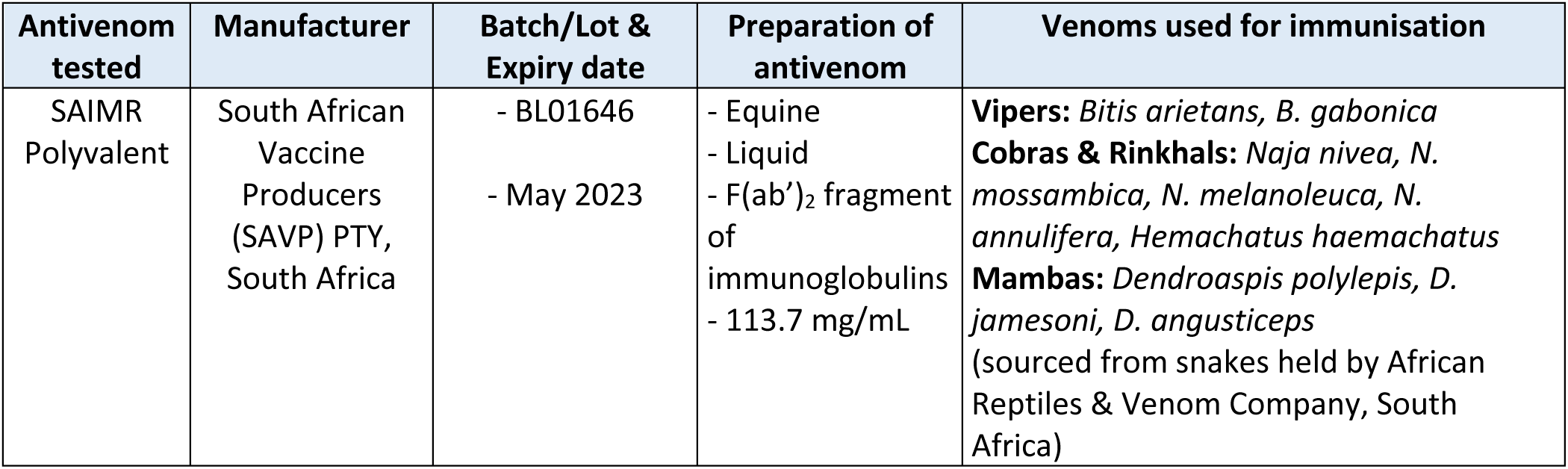
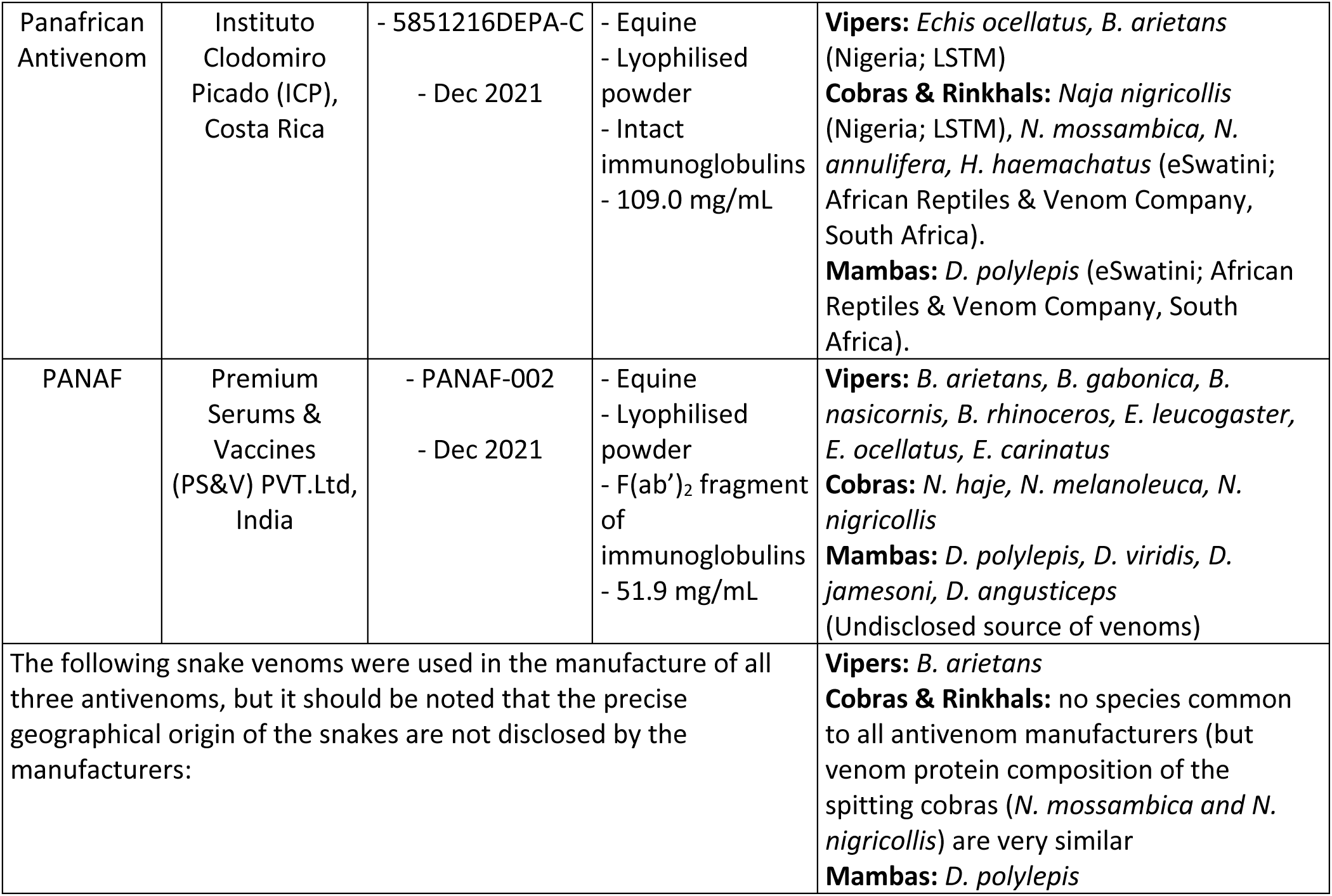
Details of the antivenom products examined in this study.

| Antivenom tested | Manufacturer | Batch/Lot & Expiry date | Preparation of antivenom | Venoms used for immunisation |
| --- | --- | --- | --- | --- |
| SAIMR Polyvalent | South African Vaccine Producers (SAVP) PTY, South Africa | - BL01646<br>- May 2023 | - Equine<br>- Liquid<br>- F(ab')<sub>2</sub> fragment of immunoglobulins<br>- 113.7 mg/mL | **Vipers:** *Bitis arietans*, *B. gabonica*<br>**Cobras & Rinkhals:** *Naja nivea*, *N. mossambica*, *N. melanoleuca*, *N. annulifera*, *Hemachatus haemachatus*<br>**Mambas:** *Dendroaspis polylepis*, *D. jamesoni*, *D. angusticeps*<br>(sourced from snakes held by African Reptiles & Venom Company, South Africa) |
| Panafrican Antivenom | Instituto Clodomiro Picado (ICP), Costa Rica | - 5851216DEPA-C<br>- Dec 2021 | - Equine<br>- Lyophilised powder<br>- Intact immunoglobulins<br>- 109.0 mg/mL | **Vipers:** *Echis ocellatus*, *B. arietans* (Nigeria; LSTM)<br>**Cobras & Rinkhals:** *Naja nigricollis* (Nigeria; LSTM), *N. mossambica*, *N. annulifera*, *H. haemachatus* (eSwatini; African Reptiles & Venom Company, South Africa).<br>**Mambas:** *D. polylepis* (eSwatini; African Reptiles & Venom Company, South Africa). |
| PANAF | Premium Serums & Vaccines (PS&V) PVT.Ltd, India | - PANAF-002<br>- Dec 2021 | - Equine<br>- Lyophilised powder<br>- F(ab')<sub>2</sub> fragment of immunoglobulins<br>- 51.9 mg/mL | **Vipers:** *B. arietans*, *B. gabonica*, *B. nasicornis*, *B. rhinoceros*, *E. leucogaster*, *E. ocellatus*, *E. carinatus*<br>**Cobras:** *N. haje*, *N. melanoleuca*, *N. nigricollis*<br>**Mambas:** *D. polylepis*, *D. viridis*, *D. jamesoni*, *D. angusticeps* (Undisclosed source of venoms) |
| The following snake venoms were used in the manufacture of all three antivenoms, but it should be noted that the precise geographical origin of the snakes are not disclosed by the manufacturers: |  |  |  | **Vipers:** *B. arietans*<br>**Cobras & Rinkhals:** no species common to all antivenom manufacturers (but venom protein composition of the spitting cobras (*N. mossambica* and *N. nigricollis*) are very similar<br>**Mambas:** *D. polylepis* |

Dependency on a single manufacturer risks harm to patients, especially when issues threaten availability or supply chains, as exemplified by the crisis that ensued following the cessation of FAV-Afrique and other African antivenom products (3,5). Such vulnerabilities and supply insecurities can lead to the dangerous use of inappropriate antivenoms which are not manufactured for, or effective against, envenoming by snakes in the region of interest (6). This can lead to low confidence in antivenoms, reduced market demands, drop in production and consequent increase in prices: diverse health, economic and commercial manufacturing factors that further perpetuate the cycle of antivenom inaccessibility (7). In Eswatini, the most medically-important snakes overlaps in several species found in South Africa. The supply of SAIMR Polyvalent to health facilities that need it the most has proved sufficiently challenging to warrant the establishment of the Eswatini Antivenom Foundation (EAF) (8) – a charity whose primary purpose is the purchase, storage and delivery of SAIMR Polyvalent to hospitals and health centres, and provision of clinical-use guidelines and training (8,9). Like other countries/agencies that have historically imported this valuable clinical product, the supply of SAIMR Polyvalent to Eswatini has become increasingly and disturbingly intermittent. Recently EAF has been unable to secure a supply of this antivenom for several months. Consequently, there is a clear need in Eswatini and other southern African countries to identify alternate antivenoms that possess the requisite clinical efficacy.

Clinical trials for new antivenoms are critical to determine their optimal initial dose, effectiveness and safety, but such trials pose significant challenges including very high cost, complex logistics, difficulty in confirming the species of snake, and heterogeneity in clinical presentations even amongst patients bitten by the same species of snake. These funding and other challenges limit efforts by the research community to conduct these urgently needed trials.

There are no agreed protocols for less expensive clinical observation studies and we are therefore reliant upon results of preclinical murine assays measuring the comparable (i) toxicity of venoms and (ii) efficacy of antivenoms. We have previously conducted these antivenom-efficacy assays to address snakebite-treatment vulnerabilities in Nigeria (10,11) and Kenya (2). To assist the research and medical community, we recently compiled all relevant published preclinical data on African snake venoms and antivenoms deployed in sSA (4). This publication was designed to link preclinical data to an important review of antivenom clinical trials performed in sSA (3), which identified that there are numerous antivenoms currently used in sSA for which no publicly available data on preclinical or clinical efficacy exists (3,4). To address this issue, the World Health Organization (WHO) have instituted an independent risk-benefit assessment of antivenom products suitable for use in sSA to underpin a subsequent prequalification scheme to support the procurement of suitable antivenom products (12). Antivenom and snakebite stakeholders are awaiting the results of this assessment, which includes data on the efficacy, stability, purity, production volumes of the products.

In the meantime, the EAF sought partners to prepare Eswatini for antivenom clinical trials necessary to identify suitable antivenom products for use in the country. With grant funding from Wellcome, EAF joined the African Snakebite Research Group (ASRG) (13) in 2019 and hosted the Eswatini Snakebite Research & Intervention Centre (E-SRIC), enabling E-SRIC to engage with and benefit from similar research and intervention objectives, challenges and successes of other SRICs in Kenya (14) and Nigeria (15). Greatly concerned by insecure and inadequate volumes of antivenom supply, one ASRG’s and E-SRIC’s priority research objectives was to determine whether there are existing suitably efficacious antivenoms to reduce Eswatini’s clinical dependency upon SAIMR Polyvalent.

Here, we preclinically examined the neutralising efficacy of SAIMR Polyvalent and two ‘test’ antivenoms against the lethal and tissue-destructive effects of venoms from five medically important Eswatini snake species in murine models of snake envenoming. The test antivenoms (see Table 1) chosen for this study were (i) pre-commercialisation stocks of the new ‘Panafrican’ product (manufactured by the Instituto Clodomiro Picado [ICP], Costa Rica and designed to to add efficacy against neurotoxic envenoming to the original tri-valent EchiTAbPLUS-ICP antivenom product) and (ii) PANAF (manufactured by Premium Serums and Vaccines [PS&V], PTY, India) – an antivenom that showed impressive polyspecific neutralising ability in our previous preclinical study of venoms of medically important East African snakes (2).

We hope that the results presented here will help guide the choice of antivenom products suitable for follow-on clinical trials, and subject to satisfactory clinical performance, may provide additional treatment options and improve antivenom supply security for snakebite patients in Eswatini.

## 2. Methods

### 2.1 Venoms and Antivenoms

Venoms were extracted from *Bitis arietans, Dendroaspis polylepis, Hemachatus haemachatus, Naja anulifera* and *Naja mossambica* snakes maintained in the E-SRIC facility and which were, with government permission (the Big Game Parks authority), wild-caught in Eswatini. Venoms were frozen, lyophilised, stored and transported to Liverpool at 4 °C. Venoms were reconstituted in PBS (pH 7.4, Gibco) and stored at -80 °C just prior to use. The same batches of venoms were used throughout the study to ensure experimental consistency.

The antivenom products examined in this study are listed in Table 1. SAIMR Polyvalent antivenom (manufactured by South African Vaccine Producers Pty [SAVP], South Africa) was acquired from a pharmacy in Eswatini. Panafrican antivenom (ICP) and PANAF antivenom (PS&V) were kindly donated by the manufacturers. All three antivenoms consist of equine antibodies. Panafrican antivenom was manufactured as intact IgG, whereas SAIMR Polyvalent and PANAF were manufactured as F(ab’)_2_ fragments of IgG. SAIMR Polyvalent antivenom was used in the liquid format, as supplied. The lyophilised Panafrican Antivenom and PANAF antivenoms were respectively reconstituted in 10 mL 0.9% saline (Severn Biotech) and 10 mL sterile medical-grade water (supplied by the manufacturer). Antivenom protein concentrations listed in Table 1 were determined by measuring the A280 nm using a NanoDrop ONE (ThermoScientific), using two dilutions of antivenom and read in duplicate. To ensure instrument accuracy, equine IgG standards were run in parallel on the NanoDrop, and a commercial BCA assay (Pierce BCA Protein Assay Kit, ThermoFisher Scientific) was used to corroborate the NanoDrop results.

### 2.2 Ethical Approval

All animal experiments were performed using protocols approved by the Animal Welfare and Ethical Review Boards of the Liverpool School of Tropical Medicine and the University of Liverpool, in accordance with the UK Animal (Scientific Procedures) Act 1986. All protocols were conducted with approval from the UK Home Office under the conditions of Project Licence P24100D38 for LD_50_ and ED_50_ experiments, and Project Licence P58464F90 for MND and eMND experiments.

### 2.3 Animal Maintenance

Male CD1 mice weighing 18-20g were obtained from Charles River (UK) and acclimatised for at least one week prior to experimentation. Mice were randomly allocated into cages of five upon arrival, and these cage allocations formed subsequent treatment groups – no further randomisation was applied. Mice were housed in specific-pathogen free facilities in Techniplast GM500 cages with nesting material (Sizzlenest zigzag fibres, RAJA) Lignocell bedding (JRS, Germany) and environmental enrichment materials. Mice were housed in room conditions of 22°C temperature with 40-50% humidity and 12/12 hour light cycles. Mice had *ad lib* access to CRM irradiated food (Special Diet Services, UK) and reverse osmosis water in an automatic water system.

### 2.4 Venom toxicity profiling (LD_50_ and MND)

The amount of venom that results in 50% lethality (LD_50_) was determined using WHO-recommended protocols (16). Doses of venom were prepared with PBS (Gibco) in a total volume of 100 µL and stored on ice until intravenous administration via the tail vein using an appropriately gauged needle. The sample size of five mice per treatment group was previously identified as the minimum number of mice to produce statistically significant results. Mice were continuously monitored over the six-hour experiment for humane endpoints (seizure, external haemorrhage, loss of righting reflex, loss of grasping reflex, hind limb paralysis and hypothermia) at which point animals were immediately euthanised using rising concentrations of carbon dioxide or cervical dislocation. Observations were performed by unblinded, mixed gender experimenters. Data were collected on time to humane endpoints and the number of deaths and survivals. The LD_50_ and 95% confidence intervals were determined using Probit analysis.

The minimum dose of venom required to induce a necrotic lesion of 5 mm diameter (MND) was determined using pre-incubated doses of venom prepared in PBS to a total volume of 50 µL and injected intradermally. Prior to injection animals were briefly euthanised with 5% isofluorane to facilitate shaving at the dose site for monitoring of lesion development. Animals were monitored three times per day for 72 hours to observe general health and to monitor lesion size. After 72 hours the animals were euthanised by rising concentrations of carbon dioxide and lesions were excised from skin and measured using calipers.

### 2.5 Antivenom efficacy profiling (ED_50_ and eMND)

Antivenoms were compared for venom neutralising efficacy using ED_50_ and eMND assays. A refined version of the WHO recommended antivenom ED_50_ was used (2,17–20), in which 3 x or 5 x the LD_50_ dose was pre-incubated at 37 °C for 30 minutes with varying amounts of each antivenom, prepared in a total volume of 200 µL with PBS and the venom/antivenom mixtures injected intravenously. Due to an error, 5.3 x LD_50_ dose was used for *D. polylepis* venom (see Supp. File S1 for details). The animals were monitored and data collected and analysed as described above for the LD_50_ assays.

For eMND assays, the dose (volume) of SAIMR Polyvalent to reduce lesion size to less than 5 mm diameter was first determined, in which the venom and antivenom (in a volume of 50 µL) were preincubated for 30 minutes at 37 °C. Venom/antivenom mixtures were injected intradermally and animals were monitored as described above for the MND experiments. The effective volume of SAIMR Polyvalent was determined and the same volume of the two test antivenoms was used to compare reduction in lesion size. Data was collected and analysed as above for the MND experiments.

These assays require the use of large numbers of experimental mice that experience pain, harm and duress. We have developed experimental protocols and steps to minimise the numbers of mice used in these assays and these are described in full (Supp. File S1).

The following series of *in vitro* assays were conducted in an effort to explain the differential efficacies of the three antivenoms to neutralise the lethal and tissue-necrotic effects of the Eswatini snake venoms.

### 2.6 Titration ELISA assays to compare the anti-venom immunoglobulin (Ig) titres of each antivenom

Venom was diluted in 100 mM carbonate-bicarbonate coating buffer pH 9.6 and coated at 100 ng per well in 96-well ELISA plates (Nunc MaxiSorp, ThermoScientific) and incubated overnight at 4°C. The following day plates were washed three times with TBS with 1% Tween20 (TBS-T), blocked with 5% non-fat milk in TBS-T and incubated for 2 hours at room temperature. Plates were washed again three times with TBS-T followed by the addition of primary antibodies (neat antivenom and normal horse IgG [BioRad] as negative control) in duplicate, at an initial dilution of 1 in 100 in blocking buffer, which were five-fold serially diluted across the plate and incubated at 4°C overnight. The following day plates were washed three times with TBS-T and incubated for 2 hours at room temperature with secondary antibody horseradish peroxidase-conjugated rabbit anti-horse IgG (Sigma A6917), diluted to 1 in 1,000 in PBS. Plates were washed three times with TBS-T followed by the addition of substrate (3% 2,2’-azino-bis(3-ethylbenzthiazoline-6-sulfonic acid), Sigma) in citrate buffer pH4.0 containing 0.1% hydrogen peroxide. Plates were incubated at room temperature for 15 minutes to develop, and the signal was measured spectrophotometrically at 405 nm on LT-4500 plate reader (Labtech).

Relative avidity ELISAs were performed as described for the end-point ELISA assays, with some exceptions; primary antibodies were added at a single dilution (1 in 10,000), and after washing the primary antibody with TBS-T, the chaotropic agent ammonium thiocyanate was added at a range of concentrations (0-8 M) and incubated for 15 minutes at room temperature. The plates were washed with TBS-T and then continued to follow the same steps as described for the end-point ELISA. The reduction percentages were then calculated; data are shown in Supplemental Figure S1.

### 2.7 SDS-PAGE and immunoblotting to compare the venom protein-binding specificities of each antivenom

For SDS-PAGE, 5 µg venom in PBS was incubated at 100 °C for 5 minutes in equal volume 2 X denaturing protein loading buffer (62.5 mM Tris-Cl pH 6.8, 25% v/v glycerol, 2% SDS, 0.75% bromophenol blue) with 30% beta-mercaptoethanol, then loaded into a 4-20% Mini-Protean TGX gel (BioRad). After electrophoresis at 200V the gels were stained with Coomassie blue (50% methanol, 40% water, 10% glacial acetic acid and 0.1% Coomassie Brilliant Blue) overnight at room temperature. Gels were de-stained 2 hours at room temperature in 50% methanol, 40% deionized water, 10% glacial acetic acid and imaged on a GelDoc (Bio-Rad).

For immunoblotting, SDS-PAGE gel electrophoresis of venoms was performed as described above but using hand-cast 15% SDS-PAGE gels. Following electrophoresis, proteins in the gels were transferred to 0.2 µm nitrocellulose membranes using the TransBlot TURBO system (BioRad). Membranes were blocked in 5% non-fat milk in TBS-T on an orbital shaker overnight at 4°C on a rocker. The following day immunoblots were washed in TBS-T for 3 × 5 minutes, then incubated with neat antivenom or normal horse IgG (BioRad) diluted 1:5,000 in blocking solution for 2 hours at room temperature. Immunoblots were washed in triplicate with TBS-T as described above, then incubated for two hours at room temperature with horseradish peroxidase-conjugated rabbit anti-horse IgG (Sigma A6917) diluted 1:1800 in PBS. Immunoblots were washed again with TBS-T and developed by the addition of DAB substrate (0.05% w/v DAB with 0.024% hydrogen peroxide) by placing the membrane into the substrate for 40 seconds, before washing with deionised water and immediately photographed.

Quantitative fluorescent immunoblotting was used to compare the binding of antivenoms to venoms. To allow for accurate quantitative comparisons using normalisation to total protein content, the appropriate amount of venom to load per lane was determined using serial dilutions to identify amounts of venom that fall within the linear range, and 5 µg was determined to be within the linear range for all 5 venoms. Subsequently, 5 µg venom in 2 X denaturing protein loading buffer was incubated at 100 °C for 5 minutes, then loaded into an Any kDa Mini-Protean TGX gel (BioRad). After electrophoresis the gels were transferred to nitrocellulose membranes using the TransBlot TURBO system (BioRad). Membranes were stained and destained with Revert Total Protein Stain 700 (LI-COR Biosciences) as per manufacturers recommendations, then imaged for 2 minutes in the 700 nm channel on the Licor Odyssey Fc (LI-COR Biosciences). Signal values for each lane were exported and normalised for each venom. Membranes were blocked in 5% non-fat milk in TBS-0.1% Tween20 on an orbital shaker for 2 hours at room temperature, then incubated with 1 in 5,000 dilution of antivenom or 1 in 50 dilution of normal horse IgG (diluted in blocking solution) overnight at 4 °C. The following day membranes were washed in TBS-0.1% Tween20 for 3 × 5-minute washes, then probed with rabbit anti-horse IgG (H&L) DyLight 800 (Rockland Immunochemicals) diluted at 1 in 15,000 in 5% non-fat milk in TBS-0.1% Tween20, and incubated at room temperature for 2 hours. Membranes were washed as above and finally with 1 × 5-minute wash in TBS prior to scanning on the Odyssey Fc in the 700 nm and 800 nm channels for 2 minutes each. The 800 nm signal for each lane was exported and normalised to total protein content using the normalisation values obtained from the 700 nm signal as described above. All images were obtained using Image Studio software (Version 5.2, LI-COR Biosciences).

### 2.8 Bovine plasma clotting assay

Two-fold serial dilutions of each antivenom were made in PBS in a volume of 10 µL per well on a 384-well plate. All conditions were tested in triplicate repeats within the plate. Venom (1 mg/mL) was added at 0.5 µL per well into the respective wells, giving a final venom concentration of 500 ng/well. Plates were incubated at 37 °C for 30 minutes then cooled to room temperature. Following incubation a Multidrop Combi reagent dispenser (ThermoFisher) was used to add 20 µL of 20 mM CaCl_2_ to each well, followed immediately by 20 µL of citrated bovine plasma (BioWest, VWR). Plates were immediately read on a FLUOstar Omega plate reader (BMG Labtech) at 595 nm absorbance for 113 minutes. The assay was performed independently in technical triplicate and outliers within one run were manually identified and excluded from analyses. The area under the curve (AUC) for each venom was calculated for each replicate and a one-way ANOVA of AUC was performed to identify significant differences from the PBS control, using Prism 9 (GraphPad Software).

### 2.9 Snake venom metalloproteinase (SVMP) assay

SVMP activity was measured using the previously described fluorogenic peptide assay (18). Briefly, 1 µg of venom (in a 1 µL volume) or equal volume of PBS was added to each well in a clear, polystyrene 384-well plate (Greiner Bio-One), followed by 10 µL of antivenom (at dilutions of 1 in 4, 1 in 8 and 1 in 16) or equal volume of PBS. Venom only, antivenom only and PBS only controls were included. The SVMP substrate ES010 (BioTechne) was suspended in reaction buffer (150 mM NaCl, 50 mM Tris-HCl pH 7.5) to 6.2 mM stock. The assay plate was incubated at 37 °C for 25 minutes and then cooled to room temperature for 5 minutes before the addition of 90 µL SVMP substrate to each well (7 µM final well concentration). The plate was immediately read at excitation 320-10 nm and emission 420-10 nm with automatic gain for 103 minutes on a CLARIOstar plate reader (BMG Labtech). All conditions were performed in quadruplicate within the plate. The RFUs at 60 minutes were collected and analysed. SVMP activity was calculated for each venom, in which ‘no antivenom’ wells represented 100% activity and the change in SVMP activity in the presence of the test antivenoms was calculated as a percentage of the ‘no antivenom’ wells.

### 2.10 Phospholipase A_2_ (PLA_2_) activity assay

PLA_2_ activity was measured using an EnzCheck Phospholipase A2 assay kit (E10217, ThermoFisher Scientific), as previously described (21). Bee venom standards were run alongside according to manufacturer’s instructions to determine specific PLA_2_ activity. Prior to analyses of the inhibitory activity of antivenoms, dilution curves of venoms only and antivenoms only were performed to (i) identify the concentrations of venom for each snake species that fall within the linear range of the activity curve, and (ii) identify whether the addition of antivenom causes assay interference. For data analysis, buffer only well values were subtracted from all other values. From these results (shown in Supplemental Figure S2), the optimal venom amounts were determined as 2 µg for *B. arietans, D. polylepis* and *N. annulifera*, 0.1 µg for *H. haemachatus* and 0.05 µg for *N. mossambica*, and antivenom volumes of less than 1.25 µL per well did not interfere with the assay readout. Using these venom amounts, the specific PLA_2_ activity was calculated from a standard curve using a bee venom positive control as per the manufacturers recommendations.

Two-fold dilutions of antivenom (diluted in PBS in a 12.5 µL volume) were incubated with the pre-defined amount of venom (in 2.5 µL volume) at 37 °C for 30 minutes, cooled to room temperature and added to a clear, polystyrene 384-well plate (Greiner BioOne). 12.5 µL PLA_2_ substrate (reconstituted as per manufacturers instructions) was added to each well, the plate was then incubated in the dark at room temperature for 10 minutes and then read at excitation 485 nm and emission 520 nm with gain set to 50% of the highest amount of bee venom control, using a FLUOstar plate reader (BMG Labtech). RFU measurements were converted to percentage of PLA_2_ activity, where the venom only (no antivenom) control was 100% activity, and the data from venom plus test antivenom wells were expressed as a percentage of 100% activity. Neutralisation by varying doses of antivenom were compared to the ‘no antivenom’ control using a one-way ANOVA.

## 3. Results

### 3.1 *In vivo* neutralisation of venom bioactivity

#### 3.1.1 Venom toxicity profiling

The amount of venom that kills 50% of mice when injected intravenously was determined for each venom, as shown in Table 2. The five venoms tested demonstrated a wide range in lethal dose. Venoms from *D. polylepis* and *N. mossambica* demonstrated the greatest lethal toxicity whereas *H. haemachatus, N. annulifera* and *B. arietans* showed weaker toxicity in the intravenous murine model.

**Table 2.**
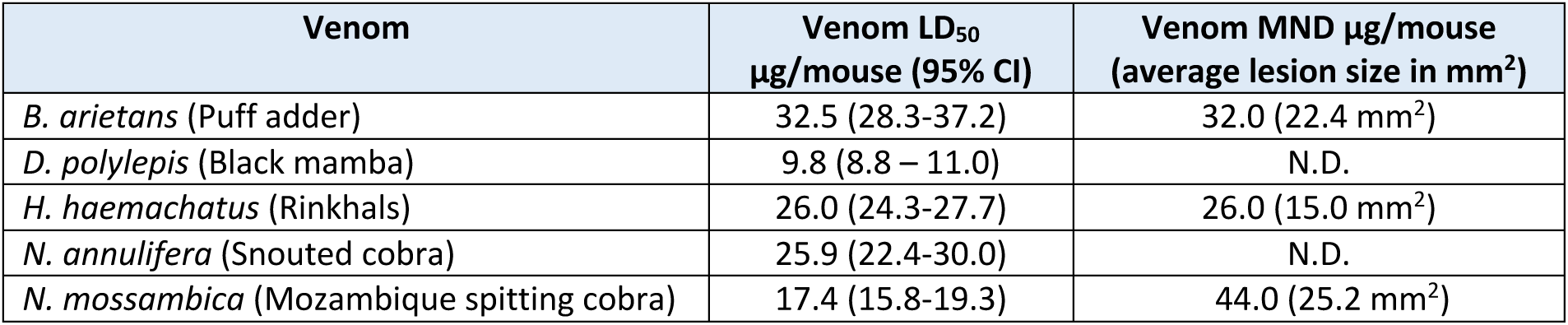
Lethal toxicity of venoms (Venom LD_50_) and necrotic toxicity (Minimum Necrotic Dose; MND) from Eswatini snake venoms in a murine model. N.D. = not determined.

The amount of venom to cause a tissue-destructive lesion was determined for the three venoms reported to cause necrosis clinically (*B. arietans, H. haemachatus* and *N. mossambica*), is shown in Table 2. Due to overwhelming neurotoxicity induced by *H. haemachatus* venom at higher doses, the maximum lesion size obtained was lower than a typical MND (5mm x 5 mm). Prior preclinical research reported that 60 µg of Eswatini *H. haemachatus* venom did not result in a dermonecrotic lesion in venom-injected mice – we are unable to account for this difference to our results (22).

#### 3.1.2 Antivenom efficacy profiling – prevention of lethal venom effects

To compare the efficacy of the antivenoms at neutralising the lethal effects of each venom, we used the WHO-recommended antivenom effective dose (ED_50_) assays, in which the amount (volume) of antivenom required to prevent venom-induced lethality in 50% of animals is determined, typically using venom concentrations of 3-5 x LD_50_.

As shown in Table 3 and Figure 1, *B. arietans* venom was most effectively neutralised by SAIMR Polyvalent (6.4 µL), whilst Panafrican required a three-fold higher dose (19.4 µL), and a drastically larger volume (47.5 µL) of PANAF was required to prevent lethality in 50% of the group. Similarly, the venom of *H. haemachatus* was most effectively neutralised by SAIMR Polyvalent (33.5 µL) and Panafrican required an almost four-fold higher dose (122.5 µL), whereas we had to use a lower challenge venom dose (3 x LD_50_) to determine the ED_50_ for PANAF, due to reaching the maximum volume limit of antivenom in the assay which this maximal volume did not confer 50% survival. Even at a reduced amount of venom, PANAF antivenom was markedly less efficacious than SAIMR Polyvalent (Figure 1).

**Table 3.** The preclinical venom-neutralising efficacy (antivenom ED_50_) of the new PANAF and Panafrican antivenoms compared to SAIMR Polyvalent in the pre-incubation model of envenoming. The assays utilised a venom challenge dose of 5 venom LD_50_s (Table 2), except when it was necessary to reduce the challenge dose 3 venom LD_50_s (indicated by red text) – to avoid overwhelming lethality. Results are reported as ED_50_ in (i) volume of antivenom and (ii) as µL of antivenom per mg of venom. The latter normalises reporting to the amount of venom used, and thus differences in number of venom LD_50_s used are compensated for.

| Snake Venom | Venom-neutralising efficacy ED<sub>50</sub> reported as |  |  |
| --- | --- | --- | --- |
| | (i) $\mu\text{L}$ per mouse (95% CI) | | |
| | (ii) $\mu\text{L}$ per mg venom | | |
|  | SAIMR Polyvalent (SAVP) | Panafrican (ICP) | PANAF (PS&V) |
| *B. arietans* (Puff adder) | 6.4 $\mu\text{L}$ (6.1-6.6) | 19.4 (18.4-20.6) | 47.5 (46.6-48.4) |
| | 36.1 $\mu\text{L}/\text{mg}$ | 109.7 $\mu\text{L}/\text{mg}$ | 268.0 $\mu\text{L}/\text{mg}$ |
| *D. polylepis* (Black mamba) | 22.1 (21.2-22.9) | 79.8 (76.3-83.6) | 56.1 (53.8-58.6) |
| | 424.6 $\mu\text{L}/\text{mg}$ | 1535.4 $\mu\text{L}/\text{mg}$ | 1079.2 $\mu\text{L}/\text{mg}$ |
| *H. haemachatus* (Rinkhals) | 33.5 (32.5-34.4) | 125.5 (115.6-136.4) | 52.7 (42.3-65.5) |
| | 258.3 $\mu\text{L}/\text{mg}$ | 969.5 $\mu\text{L}/\text{mg}$ | 677.7 $\mu\text{L}/\text{mg}$ |
| *N. annulifera* (Snouted cobra) | 77.8 (73.2-82.6) | 122.3 (117.0-127.8) | 108.4 (97.5-120.4) |
| | 600.8 $\mu\text{L}/\text{mg}$ | 1573.6 $\mu\text{L}/\text{mg}$ | 1394.6 $\mu\text{L}/\text{mg}$ |
| *N. mossambica* (Mozambique spitting cobra) | 80.3 (72.3-89.1) | 82.6 (70.9-96.1) | 68.0 (64.1-71.9) |
| | 922.5 $\mu\text{L}/\text{mg}$ | 949.1 $\mu\text{L}/\text{mg}$ | 781.0 $\mu\text{L}/\text{mg}$ |

**Table 4.** The resulting lesion size after intradermal injection with one MND dose of venom preincubated with each of the antivenoms. The dose of SAIMR Polyvalent in µL was first determined, and the test antivenoms at this fixed volume were then tested to compare lesion size.

| Venom | Efficacy ($\mu\text{L}$) of each antivenom to neutralise necrotic lesion after injection with 1 x MND dose | | |
| --- | --- | --- | --- |
|  | SAIMR Polyvalent (SAVP) | Panafrican (ICP) | PANAF (PS&V) |
| *B. arietans* | 10 $\mu\text{L}$ = 1.1 mm<sup>2</sup> necrotic lesion | 10 $\mu\text{L}$ = 0 mm<sup>2</sup> necrotic lesion | 10 $\mu\text{L}$ = 3.9 mm<sup>2</sup> necrotic lesion |
| *H. haemachatus* | 15 $\mu\text{L}$ = 0 mm<sup>2</sup> necrotic lesion | 15 $\mu\text{L}$ = 0 mm<sup>2</sup> necrotic lesion | 15 $\mu\text{L}$ = 0 mm<sup>2</sup> necrotic lesion |
| *N. mossambica* | 45 $\mu\text{L}$ = 3.6 mm<sup>2</sup> necrotic lesion | 45 $\mu\text{L}$ = 0.8 mm<sup>2</sup> necrotic lesion | 45 $\mu\text{L}$ = 0.3 mm<sup>2</sup> necrotic lesion |

**Figure 1.**
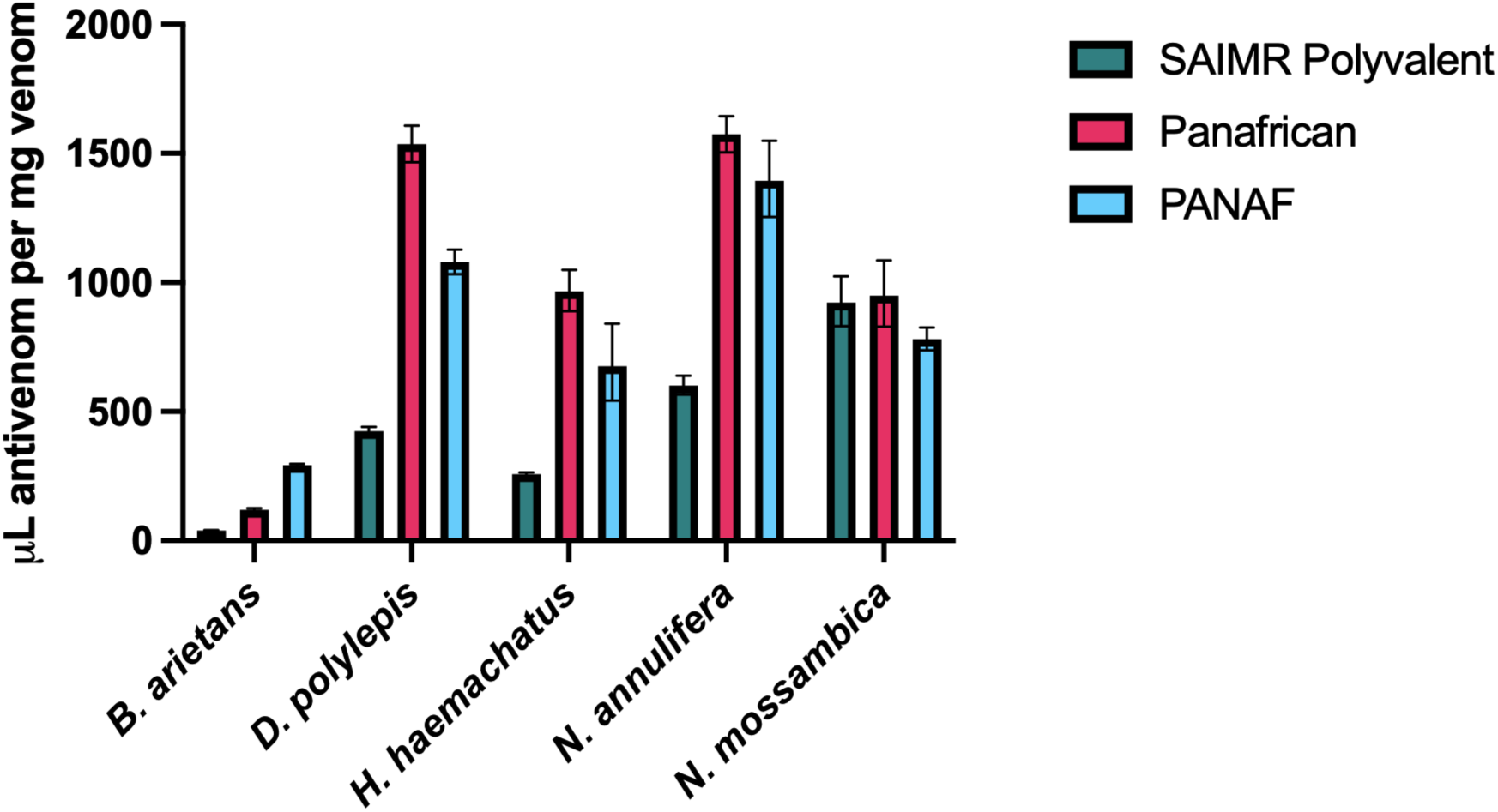
ED_50_ of the three antivenoms against each of the five Eswatini venoms, expressed as µL per mg of venom. SAIMR Polyvalent (SAVP) shown in teal, Panafrican (ICP) shown in magenta, PANAF Premium (PS&V) shown in blue. ED_50_ was determined using Probit analysis, data represents the calculated ED_50_ and error bars represent 95% confidence intervals.

Both test antivenoms were able to neutralise the lethal effects of *D. polylepis* venom, although at higher volumes than SAIMR Polyvalent, whilst for *N. annulifera* we were required to lower the venom concentration to 3 x LD_50_ as both antivenoms failed to confer 50% survival when tested at the maximum volume of antivenom with 5 x LD_50_. Even at these lower venom amounts, the volume of both test antivenoms to confer 50% survival was higher than the ED_50_ volume of SAIMR Polyvalent against 5 venom LD_50_ doses. In contrast, both test antivenoms performed equally as well or slightly better than SAIMR Polyvalent when tested against *N. mossambica* venom, with PANAF requiring the least antivenom.

As summarised in Figure 1, SAIMR Polyvalent was overall the most potent antivenom against the venoms of Eswatini snakes, with the exception of *N. mossambica* venom, which was more effectively neutralised by PANAF and Panafrican antivenom was equally as efficacious as SAIMR Polyvalent.

#### 3.1.3 Antivenom efficacy profiling – prevention of tissue-destructive venom effects

We next determined the volume of the two test antivenoms required to neutralise one MND dose of *B. arietans, H. haemachatus* and *N. mossambica* venoms. We first determined the volume of SAIMR Polyvalent to reduce the lesion size to less than 5 mm diameter. Panafrican was more effective than SAIMR Polyvalent at reducing necrotic lesion size in *B. arietans* and *N. mossambica* venoms, whereas PANAF outperformed SAIMR Polyvalent against *N. mossambica* venom but was less effective against *B. arietans* venom. All three antivenoms completely prevented dermonecrotic lesion development induced by *H. haemachatus* venom.

### 3.2 *In vitro* analyses of the interactions of antivenoms with venoms and specific venom proteins

We undertook a series of *in vitro* assays to better understand the reasons explaining the distinct efficacies of the different antivenoms to neutralise lethal and dermonecrotic effects of the various venoms. The immunological analyses examined the immunoglobulin titre and venom protein-specificity of each antivenom to each venom.

#### 3.2.1 Immunological analysis of antivenoms

##### 3.2.1.1 ELISA analysis to determine the immunoglobulin titre of each antivenom to each venom

Serial dilutions of each antivenom or normal horse IgG control were incubated with each of the five venoms in ELISA assays to compare the antivenom Ig titre to the Eswatini venoms. To represent the clinical efficacy of antivenoms, where neutralising ability is reported as volume of antivenom, we did not standardise antivenom concentrations and instead compared equivalent volumes of antivenom. The OD readings (venom-binding) of the antivenoms at the 1 in 62,500 dilution (indicated by vertical line in Figure 2 panels), were used to compare the titres of each antivenom.

**Figure 2.**
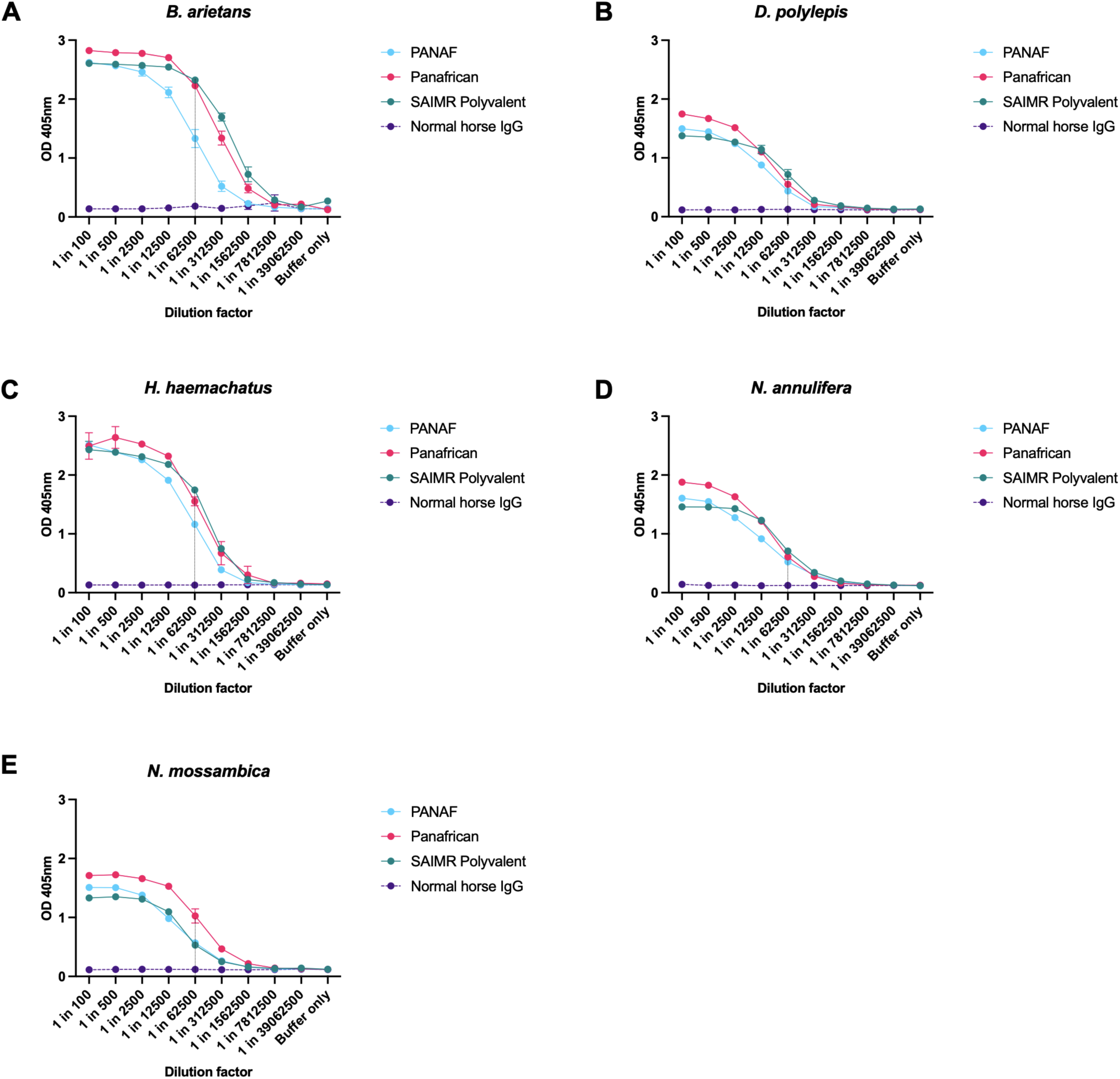
The titre of three antivenoms against five Eswatini venoms determined by end-point titration ELISA. SAIMR Polyvalent (SAVP) shown in teal, Panafrican (ICP) shown in magenta, PANAF (PS&V) shown in blue, normal horse IgG shown in purple. Panel A: *B. arietans*. Panel B: *D. polylepis*. Panel C: *H. haemachatus*. Panel D: *N. annulifera*. Panel E: *N. mossambica*. Data points represent the mean of two replicates and error bars show the standard deviation. The vertical line at the 1:62,500 dilution represents the point at which Ig titres of each antivenom were compared.

The highest overall antivenom Ig titres were against *B. arietans* venom, closely followed by *H. haemachatus* venom. At the 1:62,500 dilution ‘antivenom-comparison’ point, PANAF Premium showed a notably lower titre for *B. arietans* venom (OD_405_ = 1.33) than the SAIMR Polyvalent (OD_405_ = 2.32) and Panafrican (OD_405_ = 2.23) antivenoms. Narrower differences in Ig titre were observed for *H. haemachatus* venom, although again PANAF showed a lower titre at the (OD_405_ = 1.16) than the SAIMR Polyvalent (OD_405_ = 1.75) and Panafrican (OD_405_ = 1.55) antivenoms. Immunoglobulin titres were markedly lower for the *D. polylepis, N. annulifera* and *N. mossambica* venoms, with a highest OD_405_ reading of 1.03 (Figure 2). The three antivenoms exhibited near-equivalent Ig titres for the *D. polylepis* and *N. annulifera* venoms - differences in OD_405_ <0.3 for all antivenoms at the 1:62,500 dilution. The Panafrican antivenom exhibited the highest Ig titre (OD_405_ = 1.03) for *N. mossambica* venom – almost double the titres of the PANAF (OD_405_ = 0.57) and SAIMR Polyvalent (OD_405_ = 0.53) antivenoms. It must be noted that despite PANAF displaying the lowest Ig titres against four out of five venoms, this antivenom product contains substantially lower protein content (51.9 mg/mL; see Methods section, Table 1), which must be taken into consideration. Similarly, the antivenom with the highest protein concentration (113.7 mg/mL) SAIMR Polyvalent showed the highest Ig titres against four of the five venoms.

##### 3.2.1.2 Immunoblot analysis to determine the venom protein-specificity of each antivenom to each venom

We used immunoblot analysis to assess whether Ig titre differences between the three antivenoms were also detectable in their ability to bind distinct proteins within each venom (Figure 3). As in the ELISA analyses and for the same reason, we again used the same dilution of the antivenoms in each of the immunoblots (i.e. they were not standardised by volume not antibody content).

**Figure 3.**
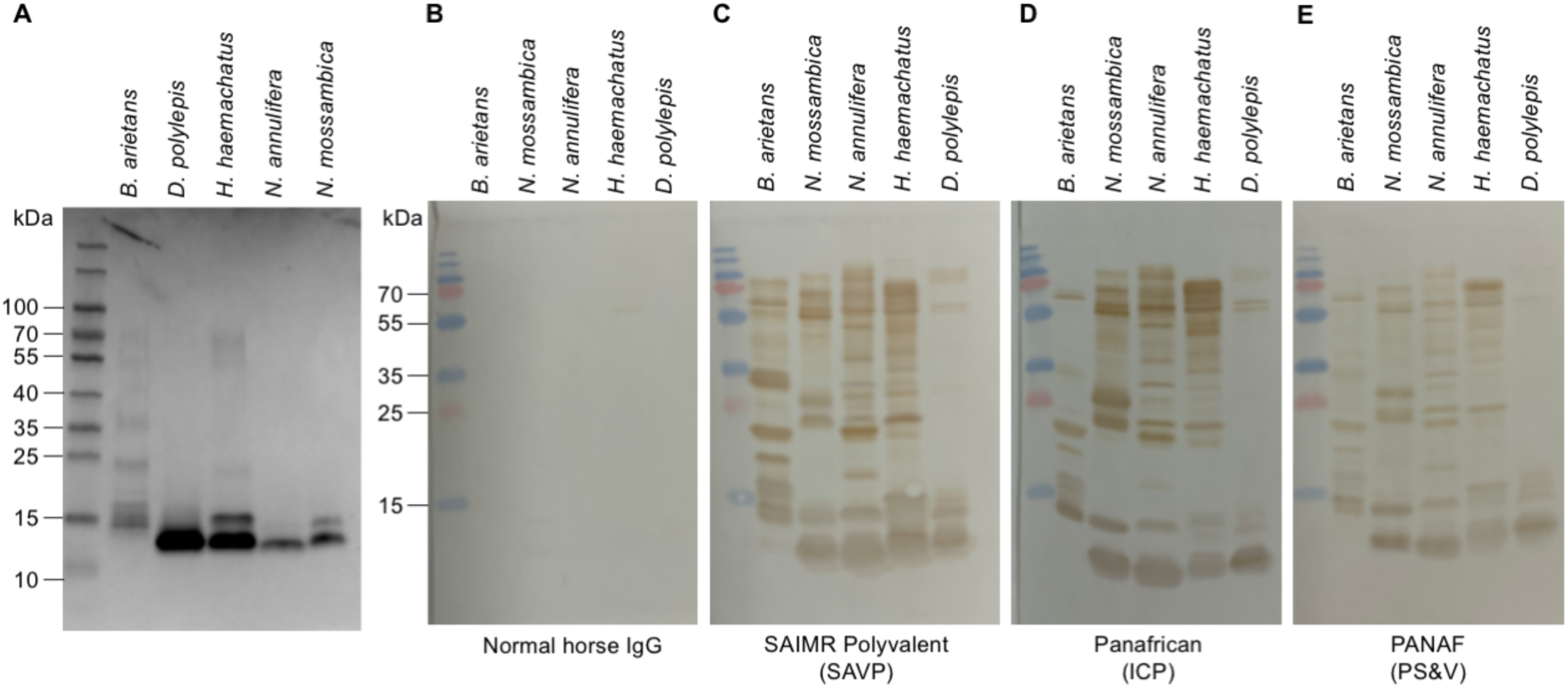
Protein profiles of the venom protein components of five Eswatini snakes. A: Venom proteins separated by SDS-PAGE under reducing conditions and stained with Coomassie blue to show all proteinaceous components. B: Western blot using normal horse IgG used as primary antibody negative control. C: Western blot using SAIMR Polyvalent as primary antibody. D: Western blot using Panafrican (ICP) antivenom as primary antibody. E: Western blot using PANAF (PS&V) antivenom as primary antibody.

All three antivenoms showed recognition of the full molecular weight range of venom proteins (depicted in Figure 3, panel A - Coomassie blue stained protein content of each venom). The PANAF antivenom showed an overall lower protein-binding intensity, however, and as per for the ELISA results, this could be attributed to the lower protein concentration of this antivenom. The most notable differences in toxin recognition by the three antivenoms was observed for *B. arietans* venom, with SAIMR Polyvalent showing stronger recognition of the higher molecular weight (MW) proteins (>25 kDa) whereas Panafrican and PANAF recognised fewer of the higher MW proteins. For the other four venoms, the toxin recognition profiles were similar across the three antivenoms, with no obvious differences in binding to specific toxins.

The immunoblot analyses above, using a chromogenic assay, failed to identify venom protein-binding distinctions that might account for the differential venom-neutralising efficacies of venom-binding Ig titres of the three antivenoms. We therefore repeated this assay using fluorescent immunoblots, which outputs quantifiable protein-binding intensities (Supplemental Figure S3). We were limited to performing these as semi-quantitative, rather than quantitative, measurements of protein binding, due to the varying formats of the antivenom Igs (F[ab’]_2_ and intact IgG), which may affect recognition by the secondary antibody thus limiting the comparisons. Aside from a more-pronounced binding of the Panafrican to the higher molecular weight components of *H. haemachatu* venom, the results using fluorescent immunoblotting did not identify any additional venom protein-binding specificities.

Altogether the serological analysis of antivenom binding to venom toxins showed similar protein binding characteristics and, with the exception of *B. arietans* venom, these data did not explain the distinct *in vivo* efficacies of the three antivenoms. Whilst the data obtained with these immunological assays is informative, it does not differentiate between recognition of toxin and non-toxin venom components and between binding and neutralising antibodies. We therefore sought to further characterise the neutralising ability of the antivenoms using *in vitro* phenotypic and toxin-specific enzymatic assays.

#### 3.2.2 *In vitro* neutralisation of venom bioactivity

#### 3.2.2.1 Plasma clotting activity

Severe anticoagulant disturbances to the normal clotting of human plasma were induced by *H. haemachatus* and *N. mossambica* venoms (Figure 4A), while venoms of *B. arietans, D. polylepis* and *N. annulifera* showed no significant alterations to plasma clotting (p > 0.05 compared to PBS control). SAIMR Polyvalent failed to neutralise the effects of *N. mossambica* venom on plasma clotting at the antivenom volumes tested, whilst Panafrican and PANAF both restored plasma clotting to ‘normal’ levels (Figure 4C). All three antivenoms effectively neutralised the anticoagulant effects of *H. haemachatus* venom to restore plasma clotting to ‘normal’ levels at the three doses tested (Figure 4B), with no significant differences between the test wells and PBS control.

**Figure 4.**
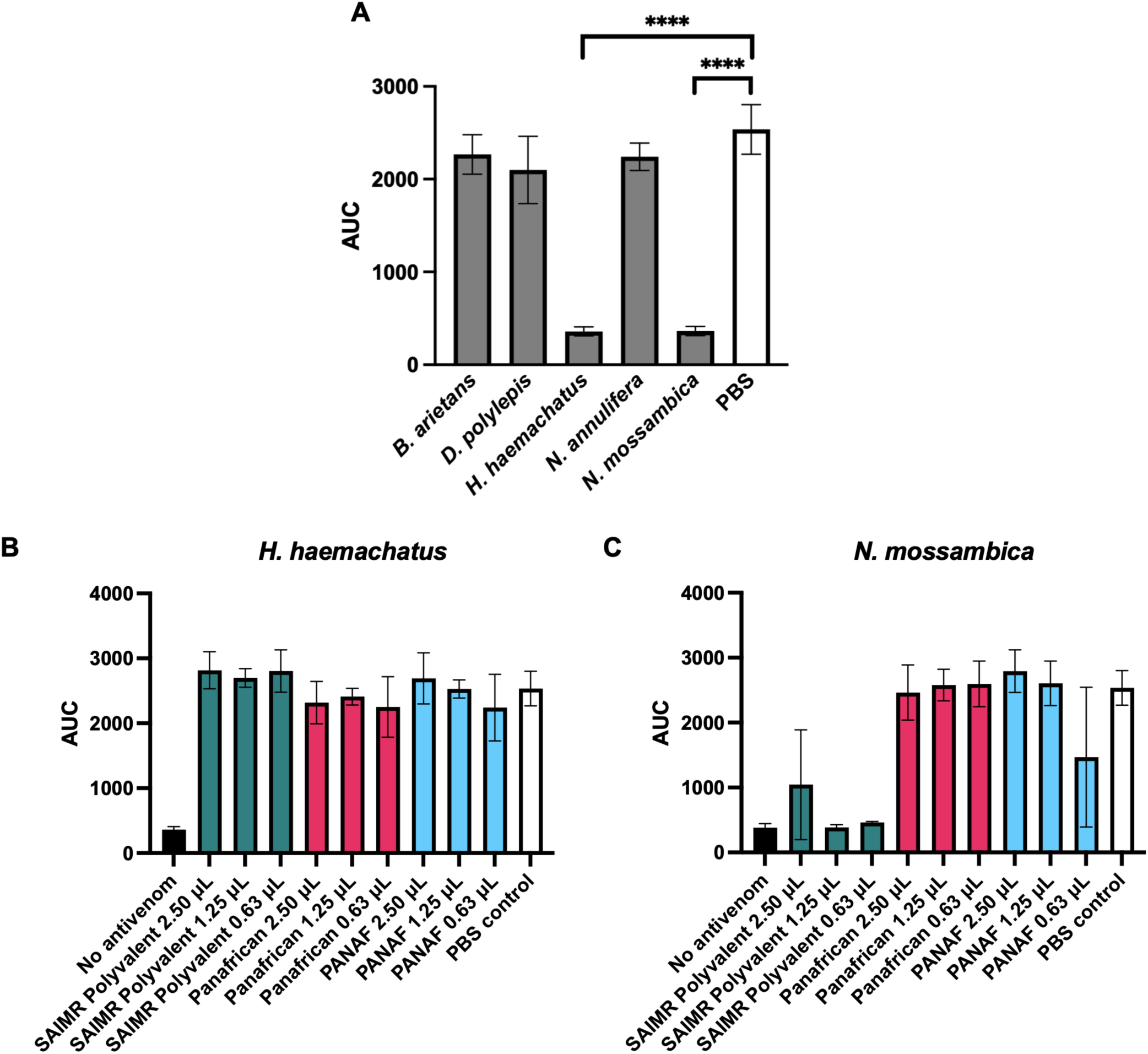
Plasma clotting activity of Eswatini venoms and their neutralisation by three antivenoms. Data show the mean area under the curve of nine replicates and error bars represent standard deviation. **** indicates p < 0.0001 by one-way ANOVA. A: Plasma clotting activity of *B. arietans, D. polylepis, H. haemachatus, N. annulifera* and *N. mossambica* in comparison to PBS normal clotting control (white bar). B: Effects of antivenoms on neutralising the anticoagulant effects of *H. haemachatus* venom. C: Effects of antivenoms on neutralising the anticoagulant effects of *N. mossambica* venom. Normal clotting is shown by ‘PBS control’ bars in white (wells contain no venom), and venom-induced clotting deficiency shown by ‘no antivenom’ control bars in black.

Although the three antivenoms showed comparable efficacy at neutralising *H. haemachatus* venom-induced anticoagulant effects *in vitro*, SAIMR Polyvalent significantly outperformed the other two antivenoms in the *in vivo* ED_50_ tests of venom lethality. The data presented here suggested that these *in vivo* differences may not be a direct consequence of clotting abnormalities.

##### 3.2.2.2 Snake venom metalloproteinase (SVMP) activity

The SVMP toxins possess potently disruptive activities against diverse mammalian cardiovascular targets (23) and we therefore compared (i) the extent to which the Eswatini venoms expressed this bioactivity *in vitro* and (ii) the SVMP-neutralising ability of the antivenoms. Strong SVMP activity was detected for *B. arietans* venom as shown in Figure 5A, whilst *N. annulifera* venom demonstrated weak but detectable SVMP activity (statistically greater than PBS control, p = 0.0004) and the remaining three venoms exhibited no significant activity (p > 0.2 for all).

**Figure 5.**
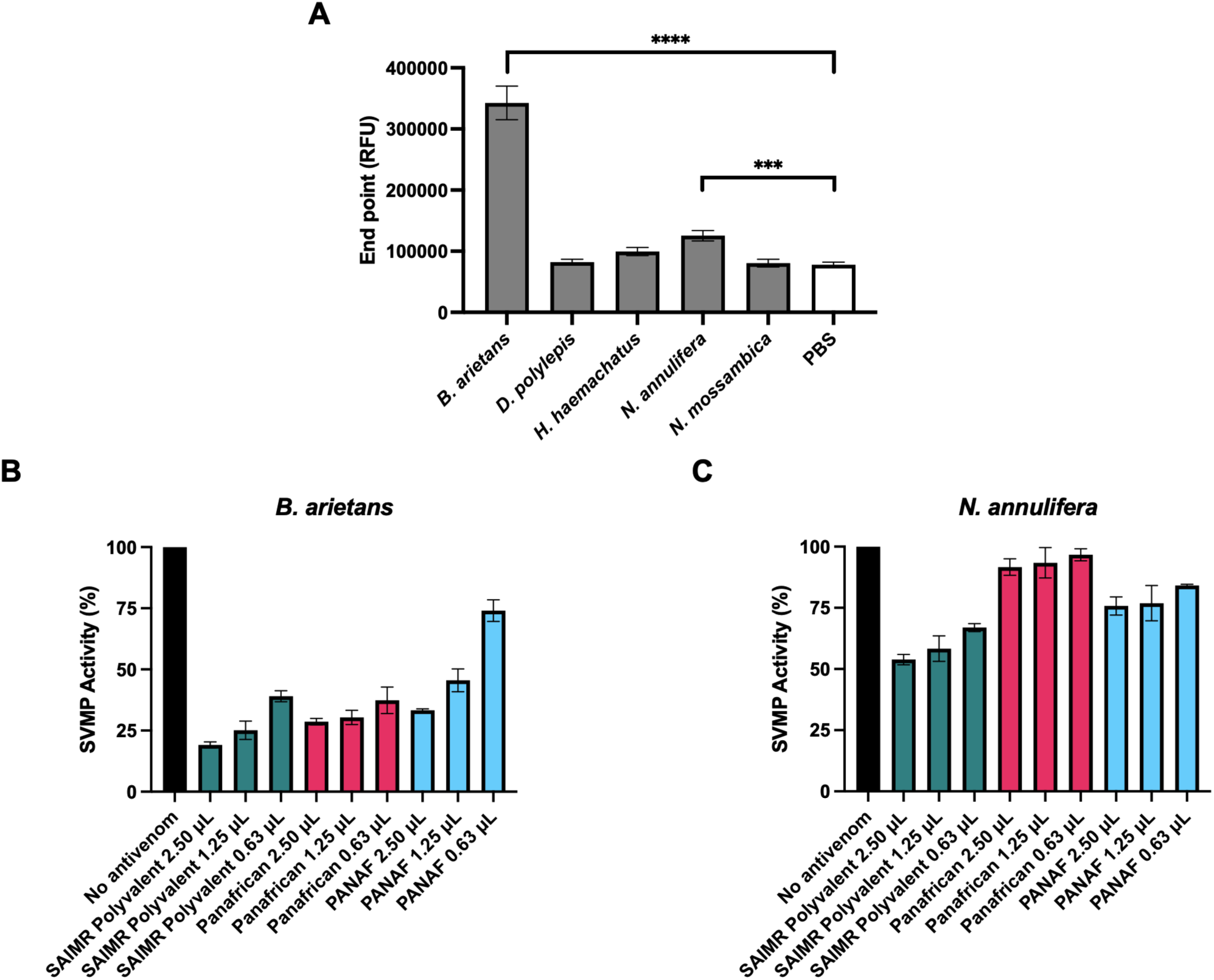
SVMP activity of Eswatini venoms and their neutralisation by three antivenoms. *** indicates p < 0.001, **** indicates p < 0.0001 by one-way ANOVA. A: SVMP activity of *B. arietans, D. polylepis, H. haemachatus, N. annulifera* and *N. mossambica* in comparison to PBS control. Data show the mean of four replicates and error bars represent standard deviation. Neutralisation of (B) *B. arietans* and (C) *N. annulifera* SVMP activity by the three antivenoms SAIMR Polyvalent (SAVP), Panafrican (ICP) and PANAF (PS&V) at different doses, in comparison to a no antivenom control (venom only) showing 100% activity. Data show the mean of four replicates and error bars represent standard deviation.

All three antivenoms significantly reduced the SVMP activity of *B. arietans* in a dose-dependent manner (p < 0.0001 for all conditions compared to no antivenom 100% activity control), with SAIMR Polyvalent demonstrating the greatest reduction in toxin activity at the highest dose (2.5 µL), as shown in Figure 5B. At the lowest dose of antivenom (0.63 µL), SAIMR Polyvalent and Panafrican showed no significant differences (p > 0.05), however PANAF showed significantly reduced levels of toxin neutralisation (p < 0.0001). The abundance of SVMP activity detected in *B. arietans* is at least partially responsible for the venom pathology observed *in vivo*, and as demonstrated here, the ranking of antivenom neutralising ability correlates well between the *in vitro* and *in vivo* results, with SAIMR Polyvalent performing best *in vitro* and *in vivo*, and PANAF performing the weakest.

All three antivenoms showed poor neutralising ability against *N. annulifera* SVMP activity (Figure 5C) and were unable to neutralise more than half of the activity. This was unexpected because SVMP activity of *N. annulifera* venom was considerably weaker than *B. arietans* venom, which was effectively neutralised by the antivenoms. As shown in Figure 5C, SAIMR Polyvalent outperformed the comparator antivenoms at all doses tested, yet only reached approximately 50% reduction in SVMP activity at the top dose tested. Panafrican antivenom only reached significant reduction in toxin activity at the top dose tested (2.5 µL).

##### 3.2.2.3 Venom Phospholipase A2 activity and neutralisation by the antivenoms

The PLA2s are a diverse and complex group of venom proteins that target a variety of mammalian cardiovascular and neurological targets (24), and we therefore compared (i) the extent to which the Eswatini venoms expressed this bioactivity *in vitro* and (ii) the PLA_2_-neutralising ability of the antivenoms. Strong enzymatic PLA_2_ activity was detected in the venoms of *H. haemachatus* and *N. mossambica* as shown in Figure 6A, whilst *B. arietans, D. polylepis* and *N. annulifera* demonstrated negligible PLA_2_ activity at the venom doses tested.

**Figure 6.**
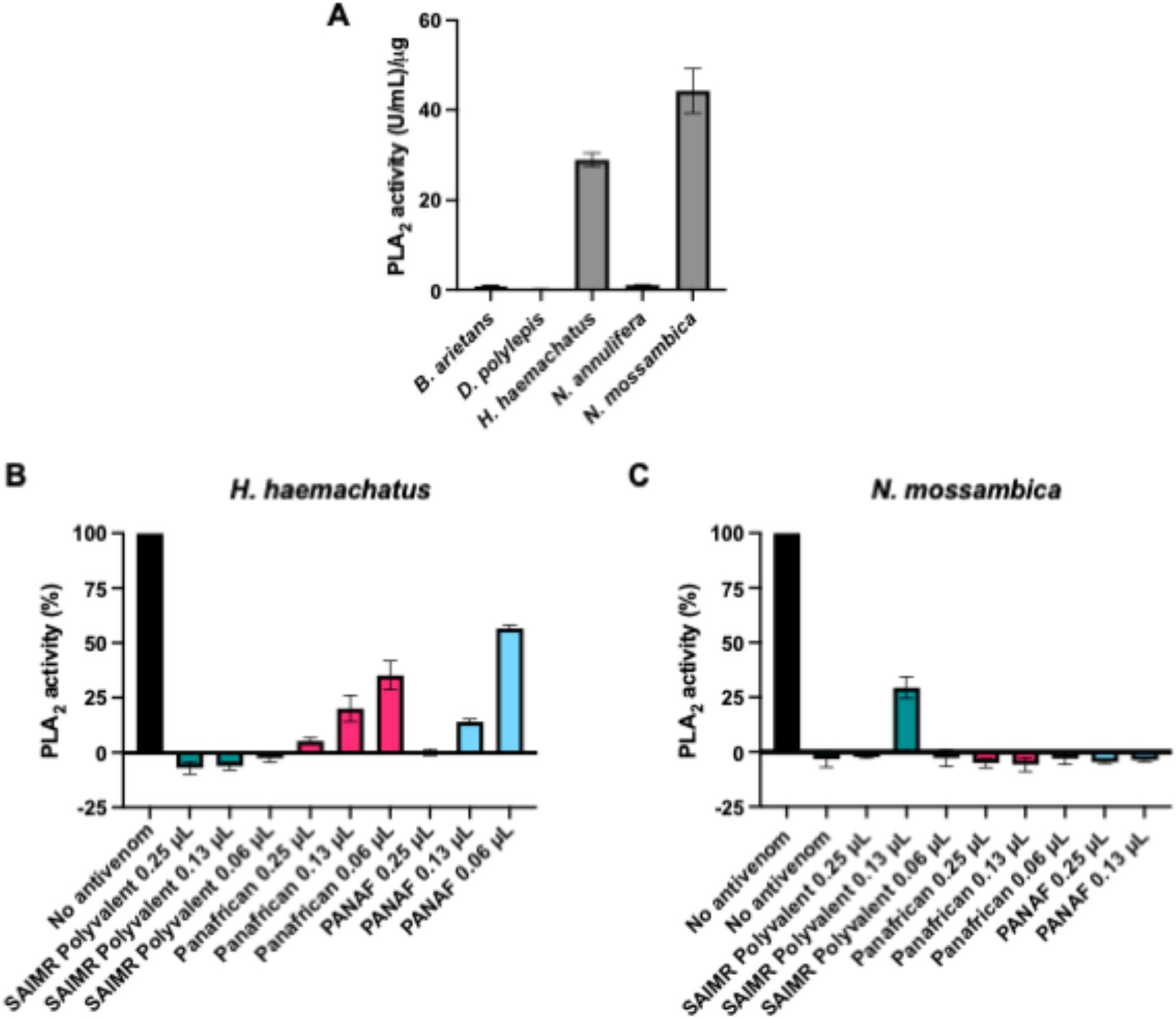
PLA_2_ activity of five venoms from Eswatini and their neutralisation by different antivenoms. A: PLA_2_ activity of *B. arietans, D. polylepis, H. haemachatus, N. annulifera* and *N. mossambica*. Data show the mean of three replicates and error bars represent standard deviation. Neutralisation of (B) *H. haemachatus* and (C) *N. annulifera* PLA_2_ activity by the three antivenom SAIMR Polyvalent (SAVP), Panafrican (ICP) and PANAF (PS&V) at different doses, in comparison to a no antivenom control showing 100% activity. Data show the mean of four replicates and error bars represent standard deviation.

All three antivenoms showed potent inhibition of both *H. haemachatus* and *N. mossambica* PLA_2_ activity, with significant neutralisation of PLA_2_ at antivenom doses as low as 0.06 µL (p < 0.0001 for each antivenom, compared to no antivenom). SAIMR Polyvalent was the most effective antivenom at neutralising *H. haemachatus* activity and was able to fully neutralise activity at the 0.06 µL dose, whereas the two other antivenoms showed lower neutralising ability at 0.13 µL and 0.06 µL doses (Figure 6B). SAIMR Polyvalent was marginally less effective than the other antivenoms in neutralising *N. mossambica* PLA_2_ activity (Figure 6C), however at doses of 0.25 µL and 0.13 µL, all three antivenoms fully neutralised the toxin activity. The ability of the antivenoms to neutralise PLA_2_ activity correlates well with the *in vivo* ED_50_ data – SAIMR Polyvalent performs best, followed by Panafrican and then PANAF. As we previously demonstrated in Figure 4 that all the antivenoms performed equally well at restoring plasma clotting, the differences in the *in vivo* antivenom ED_50_s may therefore be mediated by PLA_2_ activity, and these toxins are known to potentiate the pathological effects of other toxins found in spitting elapid venoms (25).

## 4. Discussion

The dependency of a country or region on a single antivenom product poses a high risk to the security of antivenom supply and consequent risk to victims of snake envenoming (26). There are numerous causes of such fragile antivenom markets, including manufacturing issues (inefficient and expensive production methods resulting in low production volumes (27)); logistical, technical and financial issues in procurement by, and distribution to, the communities that need the products; and low end-user product confidence (27). Access to safe and effective antivenom treatment is a pillar of the WHO strategy to halve snakebite morbidity and mortality, and the goal of delivering 3 million regionally specific treatments each year by 2030 (12). To achieve this, WHO will attempt to revitalise the antivenom market by working closely with antivenom manufacturers and end-users to improve the aforementioned deficiencies and is conducting global risk-benefit assessments of antivenoms. WHO have identified that each region should have access to at least three quality-assured antivenoms with proven effectiveness against the venoms of medically-important snakes from the region (12). To achieve this, considerable research is required to conduct preclinical and clinical testing of candidate antivenoms for each region.

The outcomes of this preclinical research collaboration between the Liverpool School of Tropical Medicine and E-SRIC, and previously with the Kenya-Snakebite Research & Interventions Centre (2) are examples of the research community responding to antivenom needs in different African countries. Here we present preclinical antivenom-efficacy data to identify effective antivenoms that can reduce Eswatini’s single-brand dependency on SAIMR Polyvalent.

Our preclinical results demonstrate that two alternative antivenom products Panafrican and PANAF, show preclinical efficacy against the venoms of medically important snakes of Eswatini. However, for four out of five venoms (the exception being *N. mossambica*), the two test antivenoms required higher doses (typically 2-3x fold more) than SAIMR Polyvalent and possessed lower snake species poly-specific efficacy. Our findings are in agreement with the previous preclinical study of Panafrican against Eswatini venoms (28). It should be noted that this was an experimental batch distinct from the pre-commercial batch of Panafrican used here. That study (28) found that SAIMR Polyvalent had a higher ability to neutralise *B. arietans, D. polylepis* and *H. haemachatus* venom lethality. Importantly, that initial batch of Panafrican antivenom failed to neutralise lethality from 3xLD_50_ of *N. annulifera* venom (27) whilst in our study the pre-commercial Panafrican was able to neutralise 3xLD_50_ of the venom. There were minor differences in the efficacy of SAIMR Polyvalent against the Eswatini venoms between our study and the previous one (27). PANAF has been previously tested for preclinical efficacy against *N. annulifera* venom sourced from Mozambique, and our ED_50_ in this study when expressed as µL/mg is similar to that obtained in the previous study, despite the different geographical origins (29). The two test antivenoms did perform as effectively or marginally better than SAIMR Polyvalent in the neutralisation of necrosis and lethality from *N. mossambica* venom. In the previous preclinical study of Panafrican, SAIMR Polyvalent was moderately more effective at neutralising *N. mossambica* lethality (28). As this snake species is responsible for the majority of snakebites in Eswatini, we believe these results justify future clinical research on the effectiveness of Panafrican and PANAF to treat envenoming in Eswatini.

This study confirms prior preclinical studies (2,28) on the potent efficacy of the SAIMR Polyvalent antivenom to neutralise the lethal effects to mice of several snake venoms, especially venoms from puff adder (*B. arietans*), black mamba (*D. polylepis*), and several spitting and non-spitting cobra (*Naja*) species, irrespective of the geographical origin of these venoms. Here, to determine the scientific reasons underpinning the extreme efficacy of the SAIMR Polyvalent antivenom, we also conducted a variety of *in vitro* assays on the bioactivity of the venoms and ability of the antivenoms to neutralise that activity – because understanding this could lead to revised antivenom-production protocols enabling the improved efficacy for all antivenoms.

Our results from the comparative ELISA and immunoblotting assays indicated that the respective antivenom titres and venom protein-specificities of Igs in the Panafrican and PANAF antivenoms were insufficiently different to account for the venom-neutralising potency of the SAIMR Polyvalent antivenom, except for *B. arietans* venom. Thus, the differences between the *in vivo* efficacies of the Panafrican and SAIMR Polyvalent against *B. arietans* venom might be explained by the observation that, in the immunoblots (Figure 3), SAIMR Polyvalent showed more intense binding to a wider molecular weight range of venom proteins than the Panafrican antivenom. Nevertheless, and as determined previously (5), the considerably higher Ig concentration of the SAIMR Polyvalent antivenom than the comparator antivenoms seemed to provide a more direct and simpler explanation than venom-binding characteristics. In this study, we deployed phenotypic and enzymatic assays of venom bioactivity because they provide toxin-focused information unattainable from the immunological assays.

The Eswatini *N. mossambica* venom completely prevented plasma clotting *in vitro* and showed strong PLA_2_ activity, as expected based upon previous observations of potent anticoagulant activity of *N. mossambica* venom (sourced from Latoxan, France) (30,31) that was attributed to PLA_2_ inhibition of Factor Xa (30). Our results indicated that Panafrican and PANAF antivenoms were more effective than SAIMR Polyvalent at neutralising the venom PLA_2_-mediated anticoagulant activity. Indeed, SAIMR Polyvalent failed to return plasma clotting to normal levels whereas the two test antivenoms both effectively neutralised this activity. These differences were reflected in the reduction of size of the tissue lesion – in which Panafrican and PANAF caused greater reductions in lesion size than SAIMR Polyvalent. That said, the three antivenoms exhibited similar efficacy in neutralising the lethal effects of *N. mossambica* venom on mice (*in vivo* ED_50_ results), though this activity may perhaps not be mediated by PLA_2_s.

The major toxin families in *H. haemachatus* venom are cytotoxic three-finger toxins (3FTx) and PLA_2_ (22,25) which produce neurotoxic and cytotoxic clinical pathologies. The Eswatini *H. haemachatus* venom exhibited an anticoagulant phenotype and strong PLA_2_ activity *in vitro*, which is in agreement with previous studies of this venom from South African specimens, which attributed the 3FTx protein family as the primary driver of anticoagulant phenotype of *H. haemachatus* venom (22). Interestingly, despite SVMPs comprising 7% of the *H. haemachatus* venom proteome, the data presented here and in previous studies show no detectable SVMP bioactivity in this venom (22). All three antivenoms performed equally well at neutralising *H. haemachatus* plasma clotting disturbances *in vitro* and preventing the development of necrotic lesions *in vivo*. Notably, SAIMR Polyvalent strongly outperformed the two test antivenoms in neutralisation of lethality assays. The discordance between *in vitro* assays of plasma clotting and *in vivo* neutralisation of lethality may be due to several factors, including the multifactorial nature of envenoming (role of other non-coagulopathic toxins in causing lethality), synergy with other toxins and venom proteins to cause pathology that is not detected in this assay, and the conditions used in this assay that do not account for phospholipids that are required to more accurately mimic physiological conditions (32).

The Eswatini *B. arietans* venom showed strong SVMP activity, negligible PLA_2_ activity and negligible effects on plasma clotting *in vitro*. This profile is similar to previous reports on the venom of *B. arietans* from Nigeria, which demonstrated strong SVMP and serine protease activity with no procoagulant effects on plasma clotting (19). Anticoagulant activities have been detected in nanofractionated *B. arietans* venom from Nigeria (33) and whole venom from South Africa (34). Here, SVMP bioactivity was inhibited by similar doses of the SAIMR Polyvalent and Panafrican antivenoms, and higher doses (volumes) of the PANAF antivenom were required to achieve similar SVMP-inhibition levels. Correspondingly, more PANAF antivenom was required to neutralise venom lethality *in vivo*. However, differential antivenom neutralisation of SVMP bioactivity is not the sole explanation of *in vivo* efficacy because the antivenom ED_50_ of the Panafrican antivenom was three-fold higher than that of SAIMR Polyvalent (Table 3) – the more intense and comprehensive *B. arietans* venom-binding of the SAIMR Polyvalent than Panafrican antivenom may be linked to the distinct *in vivo* venom-neutralising efficacies of these antivenoms (see above and Figure 3).

The Eswatini *N. annulifera* venom exhibited negligible PLA_2_ activity, similar to previous reports on toxin activity and venom proteome abundance (29,35). In contrast to the previous report on *N. annulifera* venom from South Africa (35), we detected no anticoagulant effects of the venom at the venom doses tested here. Consistent with previous reports from South African samples (35) and with recent venom proteome data identifying that SVMPs comprise 11.2% of *N. annulifera* venom from Mozambique (29), we detected SVMP activity in the Eswatini *N. annulifera* venom. The *in vitro* SVMP activity of this venom was, unexpectedly, not effectively neutralised by any of the antivenoms at the doses tested – with all three antivenoms failing to reduce SMVP activity by half. The ELISA and immunoblots also show equivalent immune-binding to the *N. annulifera* venom by all three antivenoms – but these bioactivity and immunological results do not explain the much greater *in vivo* neutralisation of *N. annulifera* venom-induced lethality by the SAIMR Polyvalent antivenom.

Highlighting the lack of *in vitro* neutralisation assays for neurotoxic bioactivity, none of the *in vitro* assays used in this study detected toxin activity of *D. polylepis* venom, which is mostly comprised of neurotoxic Kunitz-type serine protease inhibitors and 3FTx (21,36). Immunological data in this study did not show any differences in the recognition of *D. polylepis* venom, yet the two test antivenoms were markedly less effective at neutralising the venom lethality *in vivo*. Recently developed *in vitro* assays that measure neurotoxin binding to isolated nicotinic acetylcholine receptors (37) are reported to show good correlation with *in vivo* neutralisation (38) and should be used in future studies to provide *in vitro* evidence for neutralisation of neurotoxins prior to *in vivo* testing. Furthermore, the development of ‘antivenomics’ to quantify the reactivity of antivenoms towards venoms provides unparalleled serological data to inform preclinical studies (39). Including the experiments described above might have helped identify antigen specificities within the Ig repertoire of the SAIMR Polyvalent antivenom that are lacking in the other antivenoms.

The limitations of this study are the absence of standardising antivenom concentrations, using gold-standard comparisons to assess neutralisation of necrosis, and non-blinding of the preclinical experimenters. We opted to perform ED_50_ assays using antivenoms as formulated by the manufacturers (as opposed to standardising them to one protein concentration) enabling us to compare the antivenoms in the form they would be used clinically. This enables practitioners to make judgements as to whether more vials of product would be needed. We did not repeat these experiments using standardised protein concentrations (to compare the gram for gram efficacy of antivenoms) because, when combined with the protein concentration information, this information can be inferred from the results presented here as volume. To assess the comparative anti-necrosis efficacy of the two test antivenoms to SAIMR Polyvalent (the gold-standard comparator) we first determined the eMND (the dose/volume required to neutralise necrosis) of the SAIMR Polyvalent antivenom for the *B. arietans, H. haemachatus* and *N. mossambica* venoms. We then compared the reduction in lesion size using that volume of the comparator antivenoms (mixed with the same amounts of venoms). This approach, rather than using full dose-finding eMND assays, provided the answer to our research question but with a considerably lower number of mice than if we had used conventional eMND assays, and was therefore the most ethical approach. Finally, investigators were not blinded to the identity of the venoms and antivenoms during the preclinical experiments because of our need to deploy venom-specific humane endpoints as a 3Rs implementation to reduce the duration of venom-induced pain, harm and distress to the mice.

In conclusion, the results from this study have (i) confirmed the high potency of the SAIMR Polyvalent antivenom to preclinically neutralise the lethal and necrotic effects induced by Eswatini’s most medically-important snake venoms, (ii) identified that the Panafrican and PANAF antivenoms possess similar efficacy as SAIMR Polyvalent antivenom against the lethal and tissue-destructive effects of *N. mossambica* venom – the most common cause of snakebite in Eswatini. However, the Panafrican and PANAF antivenoms possess inferior efficacy to SAIMR Polyvalent antivenom in neutralising the lethal effects of the other venoms tested. Our *in vitro* analyses, whilst informative, were unable to identify the venom protein specificities of the SAIMR Polyvalent antivenom that were absent in the Panafrican and PANAF antivenoms to account for their distinct *in vivo* venom-neutralising efficacies.

This data and our inferences therefrom have been reported to the Eswatini Ministry of Health. We believe they provide a clear rationale for an assessment of the safety and efficacy of these additional antivenom products in Eswatini patients of snakebite envenoming. This study has therefore contributed to a potential solution to the current dependency on a single brand of antivenom that is increasingly difficult to acquire in Eswatini, and other southern African countries solely dependent upon the SAIMR Polyvalent antivenom to treat their snakebite victims.

## Acknowledgements

We gratefully acknowledge Premium Serums and Vaccines PTY India and Instituto Clodomiro Picado Costa Rica for the provision of antivenoms used in this study. We are also grateful to the staff of the Biomedical Services Unit (University of Liverpool) for animal care, the Eswatini Big Game Parks for authorising capture and extraction of venom from the snakes, snake rescue volunteers from Eswatini who caught the snakes from the wild, the staff of the Eswatini Antivenom Foundation snake facility for snake care, Mike Perry (African Reptiles & Venom) for snake venom extractions.

## Supplemental Figures

**Fig S1.**
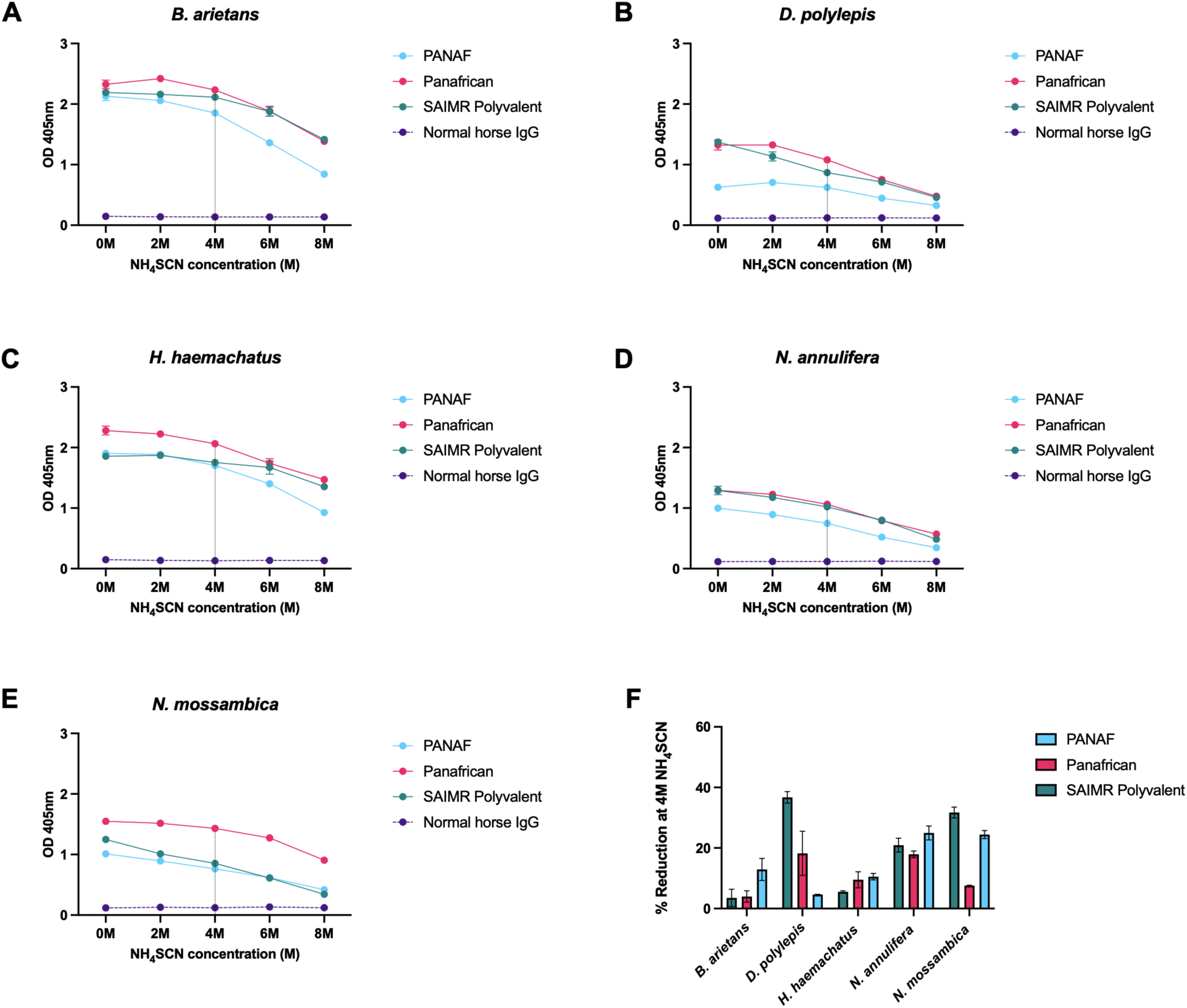
ELISA avidity

**Fig S2.**
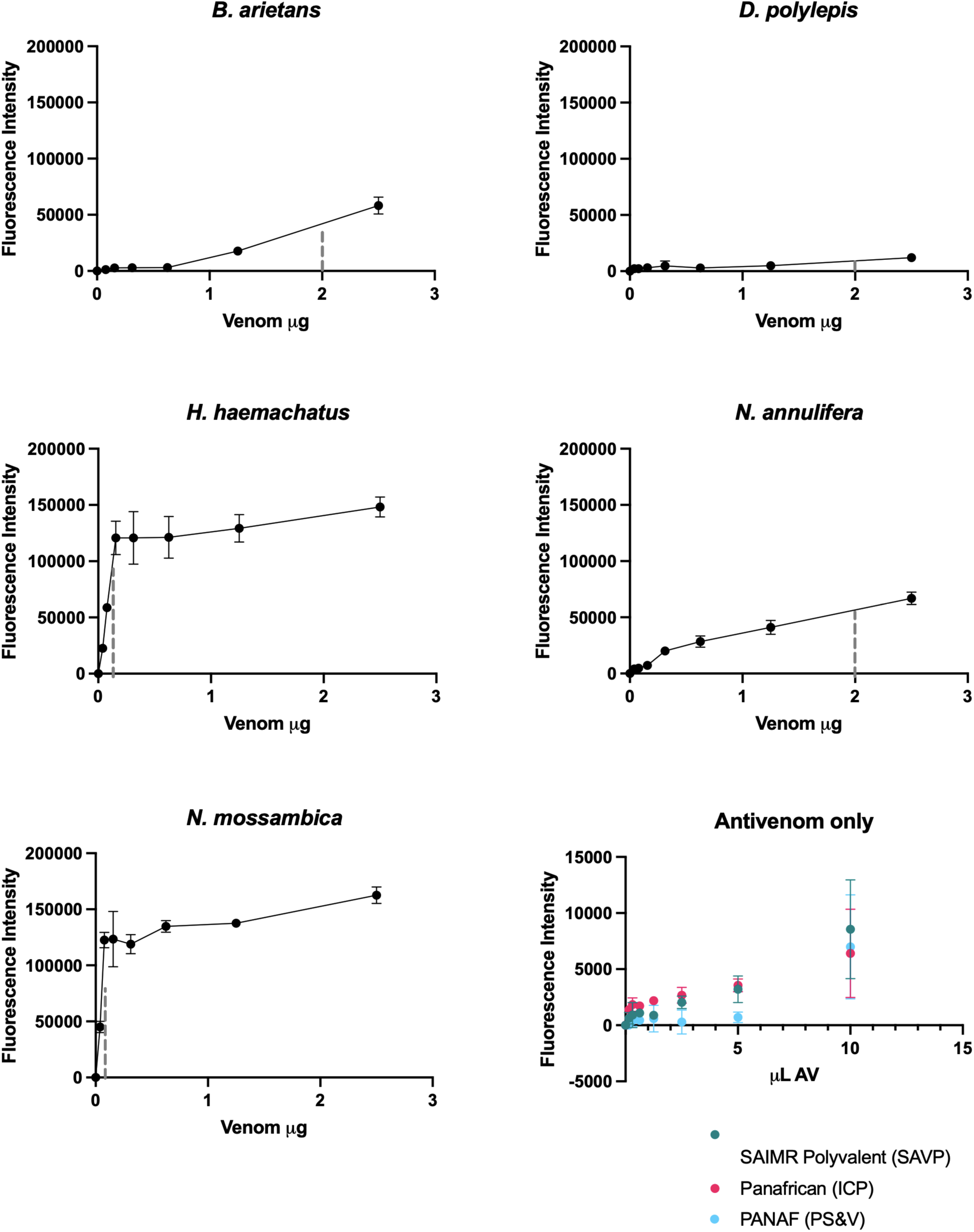
PLA_2_ pre-assay optimisation

**Fig S3.**
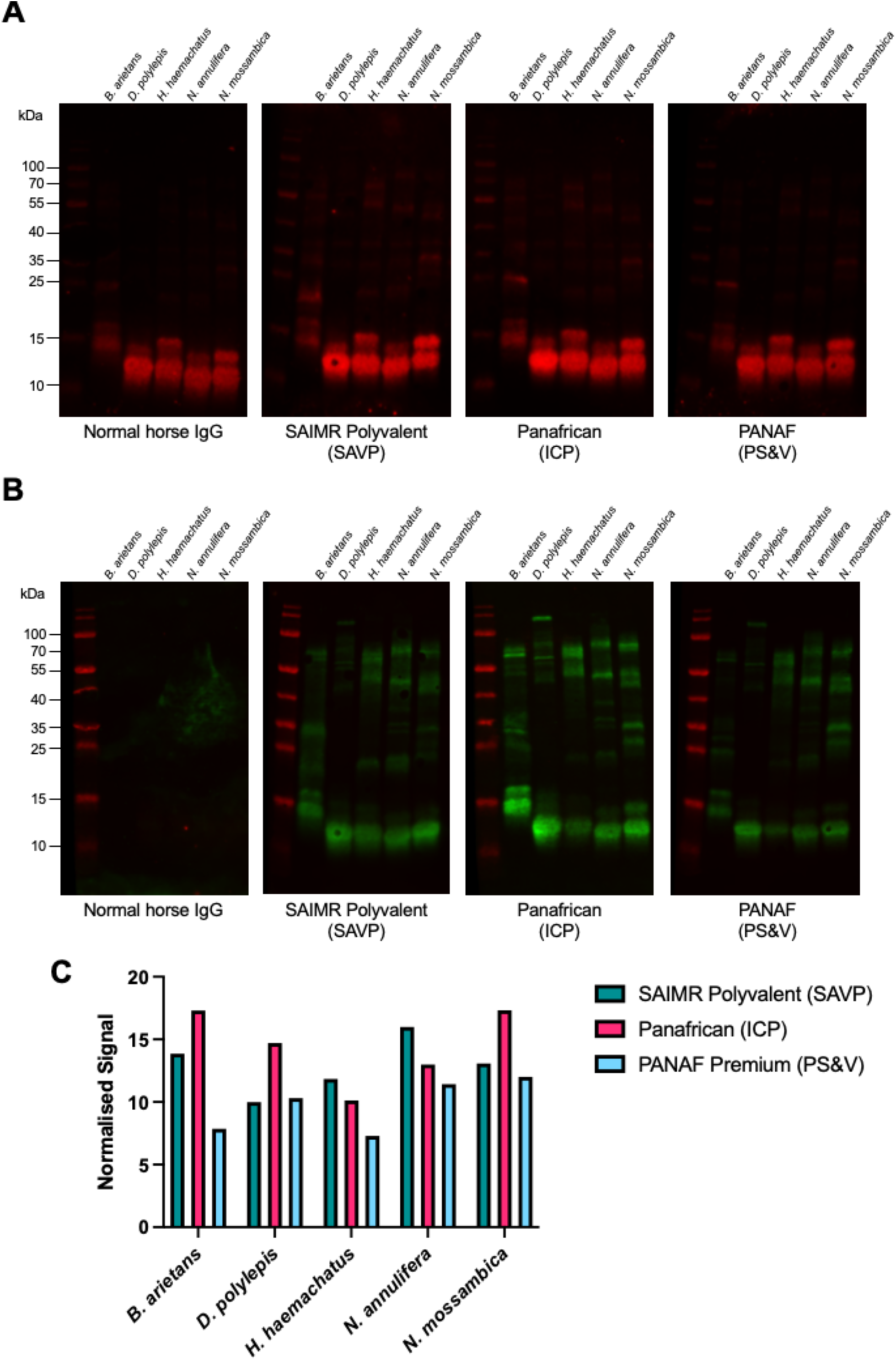
Fluorescent Western blots

## Supplemental Files

- File S1 Preclinical testing protocol
- File S2 LD_50_ and ED_50_ preclinical data
- File S3 MND measurements and eMND measurements
- File S4 ELISA endpoint raw data and ELISA avidity raw data
- File S5 Fluorescent Western blot quantification and normalisation values
- File S6 Plasma clotting assay raw data
- File S7 SVMP assay raw data
- File S8 PLA_2_ assay optimisation data
- File S9 PLA_2_ assay data

## References

1. Gutiérrez JM, Calvete JJ, Habib AG, Harrison RA, Williams DJ, Warrell DA. Snakebite envenoming. Nat Rev Dis Primer. 2017 Sep 14;3(1):1–21.

2. Harrison RA, Oluoch GO, Ainsworth S, Alsolaiss J, Bolton F, Arias AS, et al. Preclinical antivenom-efficacy testing reveals potentially disturbing deficiencies of snakebite treatment capability in East Africa. PLoS Negl Trop Dis. 2017 Oct 18;11(10):e0005969.

3. Potet J, Smith J, McIver L. Reviewing evidence of the clinical effectiveness of commercially available antivenoms in sub-Saharan Africa identifies the need for a multi-centre, multi-antivenom clinical trial. PLoS Negl Trop Dis. 2019 Jun 24;13(6):e0007551.

4. Ainsworth S, Menzies SK, Casewell NR, Harrison RA. An analysis of preclinical efficacy testing of antivenoms for sub-Saharan Africa: Inadequate independent scrutiny and poor-quality reporting are barriers to improving snakebite treatment and management. PLoS Negl Trop Dis. 2020 Aug 20;14(8):e0008579.

5. Theakston RDG, Warrell DA. Crisis in snake antivenom supply for Africa. The Lancet. 2000 Dec 16;356(9247):2104.

6. Harrison RA, Casewell NR, Ainsworth SA, Lalloo DG. The time is now: a call for action to translate recent momentum on tackling tropical snakebite into sustained benefit for victims. Trans R Soc Trop Med Hyg. 2019 Dec 1;113(12):835–8.

7. Gutiérrez JM, Warrell DA, Williams DJ, Jensen S, Brown N, Calvete JJ, et al. The Need for Full Integration of Snakebite Envenoming within a Global Strategy to Combat the Neglected Tropical Diseases: The Way Forward. PLoS Negl Trop Dis. 2013 Jun 13;7(6):e2162.

8. Antivenom Treatment for Snake Bites – Eswatini Antivenom Foundation [Internet]. [cited 2021 Dec 17]. Available from: https://eswatiniantivenom.org/

9. Robert A Harrison, David J Williams, Thea Litschka-Koen, Jonathan Pons, Patrick Joseph Tiglao, John David Commandante, et al. RSTMH Special Report on Snakebite Case reports of tropical snakebite victims illustrate the vital humanitarian role and challenges of community action groups [Internet]. 2019. Available from: https://rstmh.org/sites/rstmh/files/content/attachments/2021-04-01/RSTMH%20%E2%80%93%20Snakebite%20Report%202019%20v2.1.pdf

10. Laing GD, Lee L, Smith DC, Landon J, Theakston RDG. Experimental assessment of a new, low-cost antivenom for treatment of carpet viper (Echis ocellatus) envenoming. Toxicon. 1995 Mar 1;33(3):307–13.

11. Abubakar SB, Abubakar IS, Habib AG, Nasidi A, Durfa N, Yusuf PO, et al. Pre-clinical and preliminary dose-finding and safety studies to identify candidate antivenoms for treatment of envenoming by saw-scaled or carpet vipers (Echis ocellatus) in northern Nigeria. Toxicon. 2010 Apr 1;55(4):719–23.

12. Williams DJ, Faiz MA, Abela-Ridder B, Ainsworth S, Bulfone TC, Nickerson AD, et al. Strategy for a globally coordinated response to a priority neglected tropical disease: Snakebite envenoming. PLoS Negl Trop Dis. 2019 Feb 21;13(2):e0007059.

13. The African Snakebite Research Group [Internet]. LSTM. [cited 2022 Jan 19]. Available from: https://www.lstmed.ac.uk/research/centres-and-units/centre-for-snakebite-research/the-african-snakebite-research-group

14. KSRIC - Kenya Snakebite Research & Intervention Centre [Internet]. [cited 2022 Jan 20]. Available from: https://ksric-asrg.org/

15. NSRIC - Nigeria Snakebite Research & Intervention Centre [Internet]. [cited 2022 Jan 20]. Available from: https://nsric-asrg.org.ng/

16. World Health Organization. Guidelines for the Production, Control and Regulation of Snake Antivenom Immunoglobulins [Internet]. 2017 [cited 2022 Jan 31]. Available from: https://www.who.int/bloodproducts/AntivenomGLrevWHO_TRS_1004_web_Annex_5.pdf

17. Albulescu LO, Kazandjian T, Slagboom J, Bruyneel B, Ainsworth S, Alsolaiss J, et al. A Decoy-Receptor Approach Using Nicotinic Acetylcholine Receptor Mimics Reveals Their Potential as Novel Therapeutics Against Neurotoxic Snakebite. Front Pharmacol. 2019 Jul 30;10:848.

18. Albulescu LO, Hale MS, Ainsworth S, Alsolaiss J, Crittenden E, Calvete JJ, et al. Preclinical validation of a repurposed metal chelator as an early-intervention therapeutic for hemotoxic snakebite. Sci Transl Med [Internet]. 2020 May 6 [cited 2021 May 29];12(542). Available from: https://stm.sciencemag.org/content/12/542/eaay8314

19. Albulescu LO, Xie C, Ainsworth S, Alsolaiss J, Crittenden E, Dawson CA, et al. A therapeutic combination of two small molecule toxin inhibitors provides broad preclinical efficacy against viper snakebite. Nat Commun. 2020 Dec 15;11(1):6094.

20. Durán G, Solano G, Gómez A, Cordero D, Sánchez A, Villalta M, et al. Assessing a 6-h endpoint observation time in the lethality neutralization assay used to evaluate the preclinical efficacy of snake antivenoms. Toxicon X. 2021 Nov 1;12:100087.

21. Ainsworth S, Petras D, Engmark M, Süssmuth RD, Whiteley G, Albulescu LO, et al. The medical threat of mamba envenoming in sub-Saharan Africa revealed by genus-wide analysis of venom composition, toxicity and antivenomics profiling of available antivenoms. J Proteomics. 2018 Feb 10;172:173–89.

22. Sánchez A, Herrera M, Villalta M, Solano D, Segura Á, Lomonte B, et al. Proteomic and toxinological characterization of the venom of the South African Ringhals cobra Hemachatus haemachatus. J Proteomics. 2018 Jun 15;181:104–17.

23. Snake Venom Metalloproteinases | Charlotte A. Dawson, Stuart Ainsworth [Internet]. [cited 2022 Apr 13]. Available from: https://www.taylorfrancis.com/chapters/edit/10.1201/9780429054204-28/snake-venom-metalloproteinases-charlotte-dawson-stuart-ainsworth-laura-oana-albulescu-nicholas-casewell

24. Kini R. Excitement ahead: structure, function and mechanism of snake venom phospholipase A2 enzymes. Toxicon. 2003 Dec 1;42(8):827–40.

25. Kazandjian TD, Petras D, Robinson SD, van Thiel J, Greene HW, Arbuckle K, et al. Convergent evolution of pain-inducing defensive venom components in spitting cobras. Science. 2021 Jan 22;371(6527):386–90.

26. Potet J, Beran D, Ray N, Alcoba G, Habib AG, Iliyasu G, et al. Access to antivenoms in the developing world: A multidisciplinary analysis. Toxicon X. 2021 Nov 1;12:100086.

27. Habib AG, Musa BM, Iliyasu G, Hamza M, Kuznik A, Chippaux JP. Challenges and prospects of snake antivenom supply in sub-Saharan Africa. PLoS Negl Trop Dis. 2020 Aug 20;14(8):e0008374.

28. Sánchez A, Segura Á, Vargas M, Herrera M, Villalta M, Estrada R, et al. Expanding the neutralization scope of the EchiTAb-plus-ICP antivenom to include venoms of elapids from Southern Africa. Toxicon. 2017 Jan 1;125:59–64.

29. Tan KY, Wong KY, Tan NH, Tan CH. Quantitative proteomics of Naja annulifera (sub-Saharan snouted cobra) venom and neutralization activities of two antivenoms in Africa. Int J Biol Macromol. 2020 Sep 1;158:605–16.

30. Chowdhury A, Lewin MR, Zdenek CN, Carter R, Fry BG. The Relative Efficacy of Chemically Diverse Small-Molecule Enzyme-Inhibitors Against Anticoagulant Activities of African Spitting Cobra (Naja Species) Venoms. Front Immunol [Internet]. 2021 [cited 2022 Feb 7];12. Available from: https://www.frontiersin.org/article/10.3389/fimmu.2021.752442

31. Petras D, Sanz L, Segura Á, Herrera M, Villalta M, Solano D, et al. Snake Venomics of African Spitting Cobras: Toxin Composition and Assessment of Congeneric Cross-Reactivity of the Pan-African EchiTAb-Plus-ICP Antivenom by Antivenomics and Neutralization Approaches. J Proteome Res. 2011 Mar 4;10(3):1266–80.

32. Seneci L, Zdenek CN, Chowdhury A, Rodrigues CFB, Neri-Castro E, Bénard-Valle M, et al. A Clot Twist: Extreme Variation in Coagulotoxicity Mechanisms in Mexican Neotropical Rattlesnake Venoms. Front Immunol. 2021 Mar 11;12:612846.

33. Xie C, Albulescu LO, Bittenbinder MA, Somsen GW, Vonk FJ, Casewell NR, et al. Neutralizing Effects of Small Molecule Inhibitors and Metal Chelators on Coagulopathic Viperinae Snake Venom Toxins. Biomedicines. 2020 Sep;8(9):297.

34. Strydom MA, Bester J, Mbotwe S, Pretorius E. The effect of physiological levels of South African puff adder (Bitis arietans) snake venom on blood cells: an in vitro model. Sci Rep. 2016 Oct 24;6(1):35988.

35. Silva-de-França F, Villas-Boas IM, Serrano SM de T, Cogliati B, Chudzinski SA de A, Lopes PH, et al. Naja annulifera Snake: New insights into the venom components and pathogenesis of envenomation. PLoS Negl Trop Dis. 2019 Jan 18;13(1):e0007017.

36. Laustsen AH, Lomonte B, Lohse B, Fernández J, Gutiérrez JM. Unveiling the nature of black mamba (Dendroaspis polylepis) venom through venomics and antivenom immunoprofiling: Identification of key toxin targets for antivenom development. J Proteomics. 2015 Apr 24;119:126–42.

37. Ratanabanangkoon K, Simsiriwong P, Pruksaphon K, Tan KY, Eursakun S, Tan CH, et al. A novel in vitro potency assay of antisera against Thai Naja kaouthia based on nicotinic acetylcholine receptor binding. Sci Rep. 2017 Aug 17;7:8545.

38. Pruksaphon K, Tan KY, Tan CH, Simsiriwong P, Gutiérrez JM, Ratanabanangkoon K. An in vitro α-neurotoxin— nAChR binding assay correlates with lethality and in vivo neutralization of a large number of elapid neurotoxic snake venoms from four continents. PLoS Negl Trop Dis. 2020 Aug 28;14(8):e0008581.

39. Gutiérrez JM, Solano G, Pla D, Herrera M, Segura Á, Vargas M, et al. Preclinical Evaluation of the Efficacy of Antivenoms for Snakebite Envenoming: State-of-the-Art and Challenges Ahead. Toxins. 2017 May;9(5):163.

