## Supplemental File 1 for "Two snakebite antivenoms have potential to reduce Eswatini’s dependency upon a single, increasingly unavailable product: results of preclinical efficacy testing"

Reducing the numbers and duration of mice suffering the pain, harm and distress invoked by preclinical assays of venom-induced toxicity and antivenom efficacy

Since its ground-breaking, and World Health Organisation-adopted, 1983 publication of assays enabling comparison of venom toxicity and antivenom efficacy [1], the Liverpool School of Tropical Medicine snakebite research unit has used evidence form these assays to guide the clinical deployment of antivenoms [2, 3, 4, 5]. Nevertheless, cognisant of the pain, harm and distress these assays cause to experimental mice, we have performed research seeking alternate protocols in assays that (i) don't involve pain [the insensate hen's egg assay; 6], (ii) that reduce pain [analgesics; 7] and (iii) that reduce the number of mice required [gold standard comparison assays, dose staging protocols, 8] and reducing the duration of these severe-ranked protocols [using humane end points rather than death as a metric, 8]. Furthermore, since the vast majority of mice used in $LD_{50}$ and $ED_{50}$ experiments, irrespective of the snake venom analysed, die within 6 hours of injection, we perform these *in vivo* assays with a 6-hour duration instead of the WHO-recommended 24 hours (8).

Completion of this study required 815 mice ($LD_{50}$ assays – 225; $ED_{50}$ assays – 470; MND assays – 40; eMND assays – 80]. Where possible, we took additional steps to reduce mouse numbers whilst ensuring that the validity of the experiments was not compromised. Often this involved the requirement of the Probit statistical tool to have at least four venom or venom-antivenom dose groups of 5 mice that (i) all die, (ii) all survive and two additional groups with different survival rates.

This can be problematic. Thus, in the 100 µl Panafrican (ICP)/*N. annulifera* $ED_{50}$ experiment all the mice died quickly after injection (rising levels of $CO_2$ because of rapid onset of paralysis – one of our humane end points), except one mouse that had been mis-injected. Because paralysis had been so uniformly rapid, instead of subjecting another mouse to certain death, we entered data into Probit as 5/5 mice died.

Furthermore, in the Panafrican (ICP)/*N.mossambica* $ED_{50}$ experiment (see table), when we entered all the data from groups A-F, Probit outputted an impossible $ED_{50}$ value of over 400 µl antivenom – we presume for the reasons noted by the experimenter at the time (see RHS column). Rather than repeat this 30-mouse experiment, we found that deleting the data from mouse group B (85 µl antivenom) enabled us to acquire an $ED_{50}$ of 82.56 (72.1-94.5), and we reported this figure in the manuscript.

Finally, we report an error in calculating the $5xLD_{50}$ dose for *D. polylepis* venom. Thus, instead of multiplying 9.8 µg venom by 5 to acquire the $5xLD_{50}$ challenge dose for the $ED_{50}$ experiments, we incorrectly used a 5x 10.4 µg venom calculation. This clerical error resulted in all the $ED_{50}$ experiments being undertaken with a 5.3 venom $LD_{50}$ challenge dose rather than 5.0 $LD_{50}$ dose. Rather than repeat these experiments with another cohort of mice, we used this challenge dose in the manuscript because all the antivenoms were tested with the same venom challenge dose.

| **Experiment M283 - Data for determining ED50:** *N. mossambica* + Panafrican ICP) | | | | | |
|---|---|---|---|---|---|
| **Group #** | **Amount venom (ug/mouse)** | **Amount antivenom (ul)** | **Number Dead** | **Number Alive** | **Experimenter Notes** |
| A | 87 | 100 | 0 | 5 | Something VERY peculiar here – 65 ul should have been a near all death group. |
| B | 87 | 85 | 1 | 4 | |
| C | 87 | 75 | 2 | 3 | |
| D | 87 | 65 | 0 | 5 | |
| E | 87 | 60 | 1 | 4 | |
| F | 87 | 40 | 5 | 0 | |
| ED50 Determination | | | | Not possible in Probit because we get ED50 values above 400 ul!! | |

**References**

1. Theakston RD, Reid HA. Development of simple standard assay procedures for the characterization of snake venom. Bull World Health Organ. 1983;61(6):949-56. PMID: 6609011; PMCID: PMC2536230.

2. Theakston RD, Reid HA. Effectiveness of Zagreb antivenom against envenoming by the adder, Vipera berus. Lancet. 1976 Jul 17;2(7977):121-3. doi: 10.1016/s0140-6736(76)92846-4. PMID: 59185.

3. Abubakar SB, Abubakar IS, Habib AG, Nasidi A, Durfa N, Yusuf PO, Larnyang S, Garnvwa J, Sokomba E, Salako L, Laing GD. Pre-clinical and preliminary dose-finding and safety studies to identify candidate antivenoms for treatment of envenoming by saw-scaled or carpet vipers (Echis ocellatus) in northern Nigeria. Toxicon. 2010. April 1;55(4):719–23. doi: 10.1016/j.toxicon.2009.10.024

4. Laing GD, Yarleque A, Marcelo A, Rodriguez E, Warrell DA, Theakston RD. Preclinical testing of three South American antivenoms against the venoms of five medically-important Peruvian snake venoms. Toxicon. 2004 Jul;44(1):103-6. doi: 10.1016/j.toxicon.2004.03.020. PMID: 15225568.

5. Laing GD, Renjifo JM, Ruiz F, Harrison RA, Nasidi A, Gutierrez JM, Rowley PD, Warrell DA, Theakston RD. A new Pan African polyspecific antivenom developed in response to the antivenom crisis in Africa. Toxicon. 2003 Jul;42(1):35-41. doi: 10.1016/s0041-0101(03)00098-9. PMID: 12893059.

6. Sells PG, Ioannou P, Theakston RD. A humane alternative to the measurement of the lethal effects (LD50) of non-neurotoxic venoms using hens' eggs. Toxicon. 1998 Jul;36(7):985-91. doi: 10.1016/s0041-0101(98)00004-x. PMID: 9690791.

7. Herrera C, Bolton F, Arias AS, Harrison RA, Gutiérrez JM. Analgesic effect of morphine and tramadol in standard toxicity assays in mice injected with venom of the snake Bothrops asper. Toxicon. 2018 Nov;154:35-41. doi: 10.1016/j.toxicon.2018.09.012. Epub 2018 Sep 27. PMID: 30268394.

8. Harrison RA, Oluoch GO, Ainsworth S, Alsolaiss J, Bolton F, Arias AS, Gutiérrez JM, Rowley P, Kalya S, Ozwara H, Casewell NR. Preclinical antivenom-efficacy testing reveals potentially disturbing deficiencies of snakebite treatment capability in East Africa. PLoS Negl Trop Dis. 2017 Oct 18;11(10): e0005969. doi: 10.1371/journal.pntd.0005969. Erratum in: PLoS Negl Trop Dis. 2020 Aug 31;14(8):e0008698. PMID: 29045429;
